# TMEM70 forms oligomeric scaffolds within mitochondrial cristae promoting *in situ* assembly of mammalian ATP synthase proton channel

**DOI:** 10.1101/2020.04.03.023861

**Authors:** Hela Bahri, Jérémie Buratto, Manuel Rojo, Jim Dompierre, Bénédicte Salin, Corinne Blancard, Sylvain Cuvellier, Marie Rose, Amel Ben Ammar Elgaaied, Emmanuel Tetaud, Jean-Paul di Rago, Anne Devin, Stéphane Duvezin-Caubet

## Abstract

Mitochondrial ATP-synthesis is catalyzed by a F1Fo-ATP synthase, an enzyme of dual genetic origin enriched at the edge of cristae where it plays a key role in their structure/stability. The enzyme’s biogenesis remains poorly understood, both from a mechanistic and a compartmentalization point of view. The present study provides novel molecular insights into this process through investigations on a human protein called TMEM70 with an unclear role in the assembly of ATP synthase. A recent study has revealed the existence of physical interactions between TMEM70 and the subunit c (Su.c), a protein present in 8 identical copies forming a transmembrane oligomeric ring (c-ring) within the ATP synthase proton translocating domain (F_O_). Herein we analyzed the ATP-synthase assembly in cells lacking TMEM70, mitochondrial DNA or F1 subunits and observe a reciprocal dependence of TMEM70 and Su.c levels, regardless of the status of other ATP synthase subunits or of mitochondrial bioenergetics. Immunoprecipitation and two-dimensional blue-native/SDS-PAGE reveal that TMEM70 forms large oligomers composed of 8 TMEM70 dimers and that TMEM70 oligomers interact with Su.c not yet incorporated into ATP synthase complexes. Moreover, discrete TMEM70-Su.c complexes with increasing Su.c contents can be detected, suggesting a role for TMEM70 oligomers in the gradual assembly of the c-ring. Furthermore, we demonstrate using expansion super-resolution microscopy the specific localization of TMEM70 at the inner cristae membrane, distinct from the MICOS component MIC60. Taken together, our results show that TMEM70 oligomers provide a scaffold for c-ring assembly and that mammalian ATP synthase is assembled within inner cristae membranes.

## Introduction

In non-chlorophyllian eukaryotes, most of the ATP is produced through oxidative phosphorylation in mitochondria. In this process, the ATP synthase uses the electrochemical proton gradient generated by the respiratory chain as the driving force to synthesize ATP from ADP and Pi (1). The mitochondrial ATP synthase is a rotary motor embedded in the inner membrane (2). It is made of 16 different polypeptides in mammals of which two are synthesized inside mitochondria from mitochondrial DNA/mtDNA genes (ATP6/Su.a and ATP8/Su.A6L) (3). All others are synthesized in the cytosol from nuclear genes and imported post-translationally (See 4, 5 for reviews).

The ATP synthase subunits are organized in two distinct domains, an extrinsic matrix domain (F1) and a membrane-embedded domain (Fo) connected by a central axis and a peripheral stalk (2) (Fig. 3D). Passage of protons through the membrane domain (proton channel) involves both subunit a (Su.a) and subunit c (Su.c). This generates the rotation of a cylinder made of 8 subunits c (the c-ring) in the membrane together with the central stalk subunits of F1 (γ, δ, and ε); this ensemble forms the rotor of the enzyme. Rotation of the central stalk within the catalytic head of F1, a hetero-hexamer of three Su.Alpha and three Su.Beta, induces conformational changes that allow the binding of ADP and Pi, the formation of ATP, and its release. The peripheral stalk (made of Su.OSCP, b, d, F6) holds the catalytic head on the one side and is anchored in the membrane on the other side where it binds to Su.a, A6L, and the other subunits of the complex including supernumerary subunits (Su.f, g, e, DAPIT, 6.8PL) with no equivalent in bacterial ATP synthase.

The monomeric complexes associate in dimers via their membrane portion and those dimers assemble in rows along the edge of cristae (6–8) where the V-shape of ATP synthase dimers would favor the positive curvature of the membranes (9–13). A recent study in budding yeast has revealed that factors involved in the biogenesis of the mtDNA-encoded subunits of the ATP synthase (notably Su. c) are preferentially retrieved in cristae rather at the inner boundary membrane (14). In mammals, where Su. c is nuclear-encoded by three isogenes (ATP5G1/2/3), it is not known whether the ATP synthase is assembled near to the sites of protein import (at inner boundary membranes) or directly within cristae.

The biogenesis of the mitochondrial ATP synthase is a sophisticate process that has been investigated mainly in yeast and mammals (4, 5, 15, 16): because of their dual genetic origin, the ATP synthase subunits are synthesized in the cytosol and in the mitochondrial matrix and are then assembled in an ordered pathway to avoid accumulation of partial intermediates that could dissipate the proton gradient or hydrolyze ATP in a futile way (17, 18). The assembly of the ATP synthase is assumed to start with the separate formation of the F1 and the c-ring, that then join to form a F1-c intermediary complex. The peripheral stalk would assemble separately and next associate with the F1-c complex together with supernumerary subunits g and e (16, 19). The mtDNA-encoded subunits a and A6L would finally be added together with the supernumerary subunits 6.8PL and DAPIT.

Defects in OXPHOS complex biogenesis often lead to proteolytic clearance of unassembled subunits by mitochondrial protein quality-control systems (20, 21). For instance, defective assembly of the F1 catalytic head leads to a strong reduction in levels of all ATP synthase subunits (22, 23) except for Su.c indicating that free c-ring is a stable entity that forms separately (24–26). In the absence of Su.c the assembly of F1 can still proceed, and F1 can be found associated with peripheral stalk subunits (27). Defective assembly at later steps, for instance due to the absence of some nuclear-encoded peripheral stalk subunits, lead to the accumulation of F1-c subcomplexes (16, 19, 28, 29), and similar assemblies also comprising peripheral stalk subunits were detected in cells devoid of mtDNA-encoded subunits (27, 30–32).

Studies in yeast have identified a quite large number of specific factors required to assemble the ATP synthase that are not part of the final complex. Only two of them involved in the assembly of the Su.Beta and Su.Alpha hetero-hexamer (Atp11 and Atp12) have known mammalian functional homologues (ATPAF1 and ATPAF2; 33), indicating that significant differences in the biogenesis of ATP synthase exist between yeast and mammals. This is further supported by the existence of an additional factor (TMEM70), with no known yeast ortholog, that is required to assemble the human enzyme.

TMEM70 was linked to ATP synthase biogenesis after mutations in its gene were associated with neonatal encephalo-cardiomyopathies induced by low levels of functional ATP synthase (34). Since this first report in 2008, a number of other patients with pathogenic mutations in TMEM70 have been reported worldwide making these mutations the most frequent cause of ATP synthase deficiencies of nuclear origin (35, 36). TMEM70 is a transmembrane protein of the inner membrane that lacks specific motifs or domains allowing to infer its properties and functions. Recently, a manuscript reporting direct interactions between TMEM70 with Su.c proposed that TMEM70 could stabilize Su.c before its insertion and assembly (37).

In this study we unveil that TMEM70 localizes to cristae membranes, where it forms large homo-oligomers composed of 8 dimers. We further show that, before assembly into ATP-synthase, increasing numbers of Su.c interact with TMEM70 oligomers. We conclude that TMEM70 provide a scaffold for the assembly of the c-ring and that, in mammalian mitochondria, assembly of the c-ring and of the ATP synthase occurs within cristae.

## Results

### TMEM70 is an inner membrane protein specifically localized in cristae

Sequence analysis of TMEM70 depicts a cleavable N-terminal mitochondrial targeting sequence, two transmembrane segments and a C-terminal hydrophilic domain (Fig. 1A). The predicted sizes of the precursor and mature protein are 29 kDa and 20.7 kDa, respectively. We generated antibodies against a mixture of two distinct peptides respectively located upstream of the transmembrane segments and at the C-terminus (Fig. 1A) and we validated the reactivity of the sera towards the two peptides by ELISA (Fig. S1A). In order to further validate these antibodies, we cloned TMEM70 cDNA into a lentiviral vector independently co-expressing the DsRed fluorescent marker targeted to mitochondria, mitoDsRed (38). Transduced 143B cells were FACS-sorted according to their red fluorescence in order to obtain a homogeneous population stably expressing TMEM70 and mitoDsRed. Our antibodies allowed us to detect a protein of the expected size in total cellular extracts, which levels are increased in transduced cells expressing exogenous TMEM70 (Fig. S1B). Cell fractionation experiments (Fig. 1B) localized the detected endogenous protein in mitochondria as expected from previous studies (39–41). We obtained similar results with transduced cells in which TMEM70 is overexpressed, which ruled out mistargeting and mislocalization of exogenous TMEM70 (Fig. S2). Moreover, endogenous TMEM70 behaved like an integral membrane protein after salt or carbonate treatment (Fig. 1C), and became accessible to proteinase k after rupturing the outer membrane by osmotic swelling (Fig. 1D). Considering that the C-terminus was shown in the matrix (41) and the predicted hairpin fold of the protein that would localize the N-terminus in the same compartment (40), our results additionally suggest that the hydrophilic loop is located in the intermembrane space. Altogether, these data reinforce the topological model of TMEM70 (Fig. 1E) proposed previously (40, 41).

**Figure 1.**
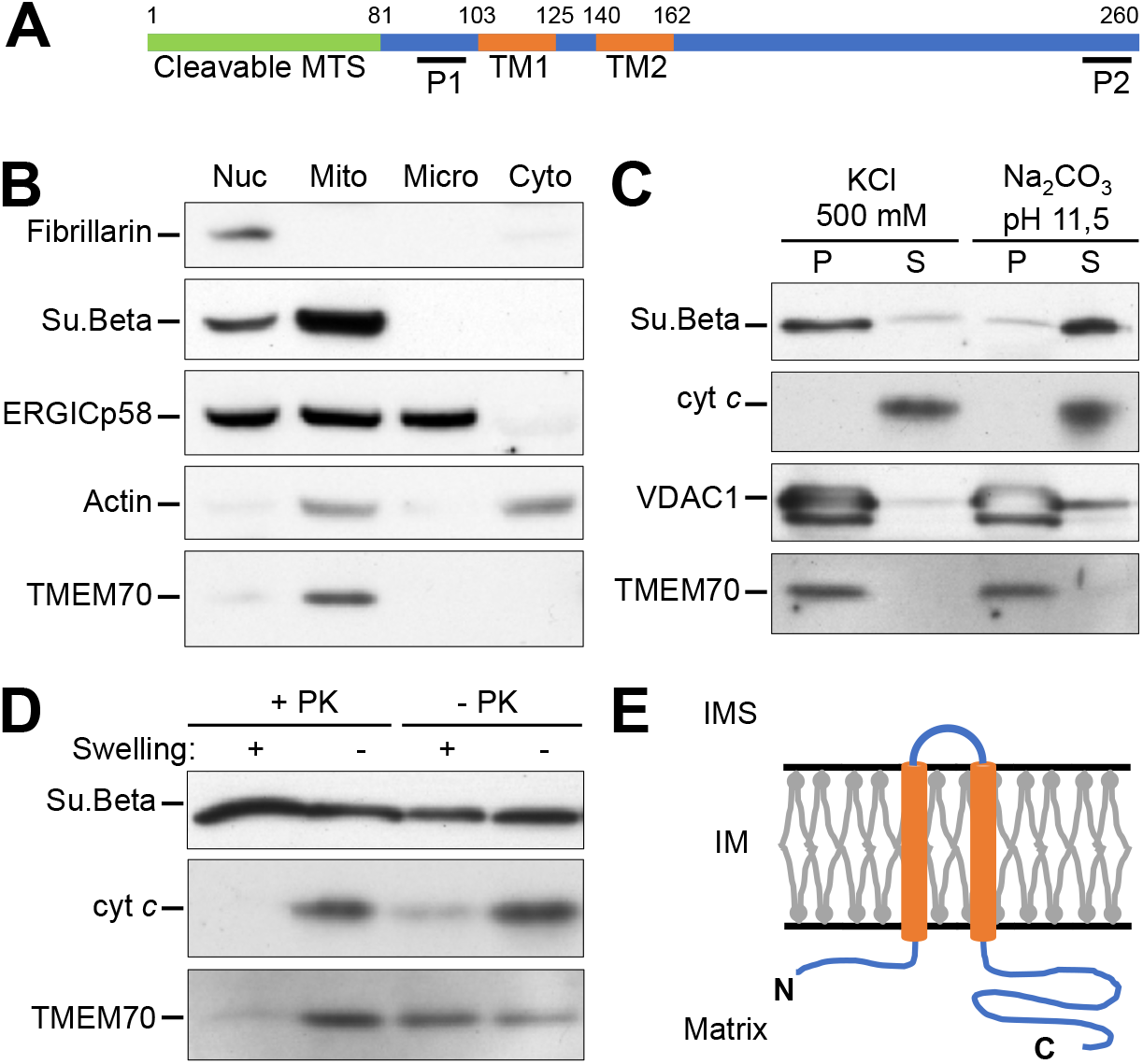
Subcellular localization of TMEM70. *A*, scheme representing the domain organization of TMEM70 with amino-acid residues numbering above. MTS: Mitochondrial Targeting Sequence. TM: Transmembrane segment. Positions of immunogenic peptides are indicated (P1, P2). *B*, 143B-rho^+^ cells were fractionated by differential centrifugation as detailed in the materials and methods section. Fractions enriched in nuclei (Nuc), mitochondria (Mito), microsomes (Micro), and cytosol were analyzed by SDS-PAGE and western blot using antibodies against indicated proteins. Protein markers of different cellular compartments/machineries were probed: Fibrillarin (nucleus), Su.Beta (mitochondria), ERGIC p58 (ER-Golgi), Actin (cytoskeleton). Endogenous TMEM70 was detected almost exclusively in the crude mitochondrial fraction as for Su.Beta. *C*, isolated mitochondria from 143B-rho^+^ cells were subjected to carbonate (Na2CO3 pH11.5) or salt (KCl 500 mM) extraction. Proteins from supernatants (Sup) and pellets were separated by SDS-PAGE and analyzed by western blot. Endogenous TMEM70 behaved like an integral membrane protein similarly to the outer membrane protein VDAC, by contrast with cytochrome *c* peripherally associated with the inner membrane and as expected released in the soluble fraction. *D*, isolated mitochondria from 143B-rho^+^ cells were subjected to a protease protection assay. Mitochondria were resuspended in hypotonic (Swelling +) or iso-osmotic buffer (Swelling -) in the presence (+PK) or absence (-PK) of Proteinase K. Protein pellets were separated by SDS-PAGE and analyzed by western blot. TMEM70, like cytochrome *c*, became accessible to proteinase k after rupturing the outer membrane by osmotic swelling while the matrix-localized ATP synthase subunit Beta remained intact, showing that the integrity of the inner membrane was preserved. *E*, topological model of TMEM70 within mitochondria. (IM) inner membrane; (IMS) intermembrane space. N- and C-termini of TMEM70 are indicated.

The ATP synthase was shown, notably in mammals, to accumulate at the inner cristae membrane (ICM) where its oligomers are involved in the shaping of this membrane (6, 7, 9, 42, 43). An as-yet unresolved question is whether the biogenesis of the ATP synthase takes place at the inner boundary membrane (IBM) that faces the outer membrane, and then moves into the ICM, or whether assembly occurs directly within the ICM. Given the involvement of TMEM70 in ATP synthase biogenesis, we addressed this question by investigating TMEM70 localization within IBM and ICM using fluorescence microscopy.

We first analyzed transduced cells expressing exogenous TMEM70 and compared its localization to that of Mitofilin/Mic60, a well characterized protein of the IBM that is a central component of the mitochondrial contact site and cristae organizing system (MICOS) localizing at cristae junctions (44). Standard confocal microscopy confirmed the localization of both proteins within mitochondria but did not allow us to reveal their submitochondrial distribution (Fig. 2A). Increasing the resolution by STED microscopy revealed a punctate pattern of both proteins throughout mitochondria. The MIC60 protein distributed to discrete puncta that appeared somewhat aligned at the mitochondria periphery, as expected (45). In contrast, TMEM70 protein distributed to smaller dots with a more homogeneous distribution and no apparent enrichment in the mitochondrial periphery.

**Figure 2.**
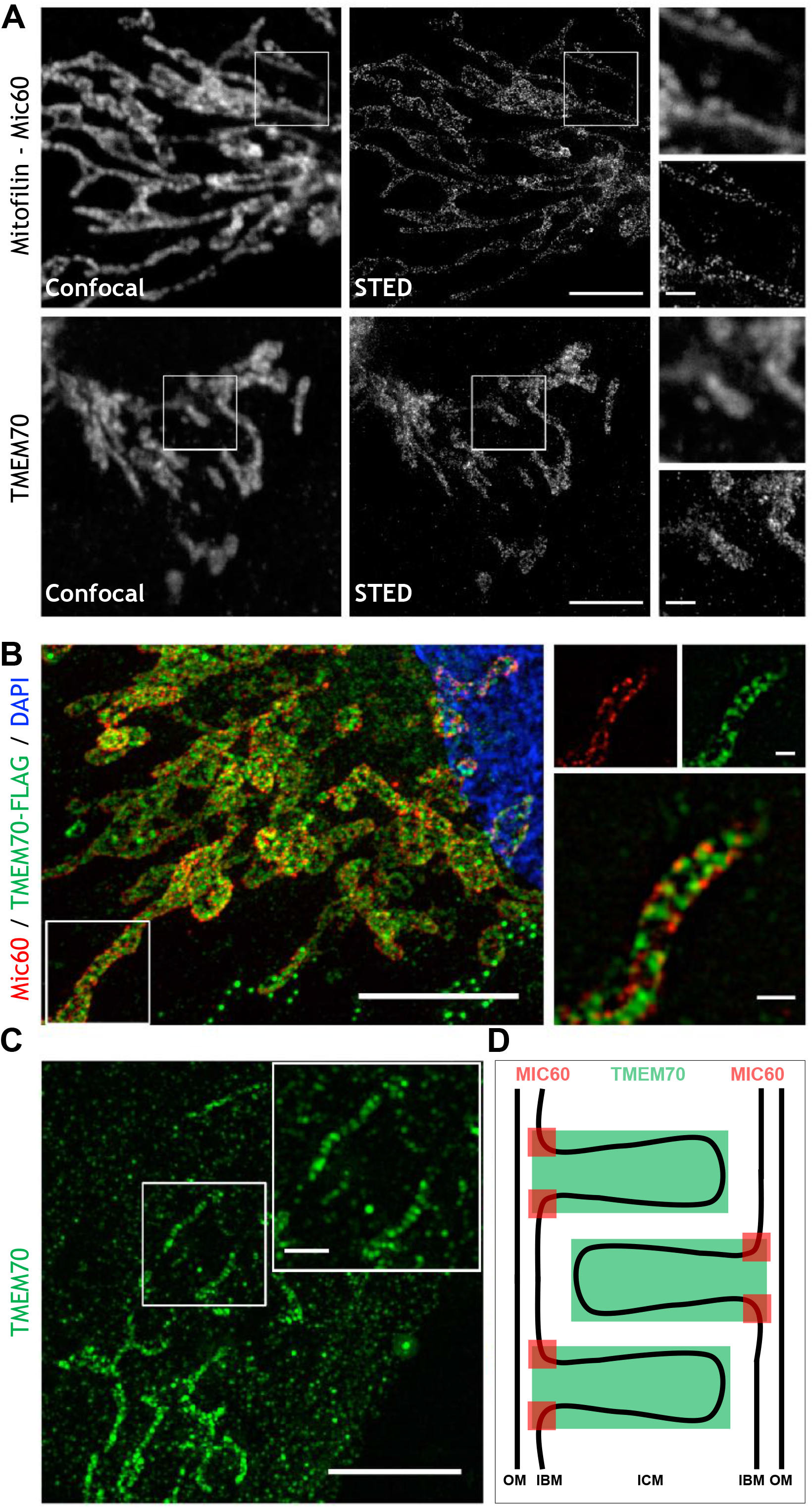
Submitochondrial localization of TMEM70 by fluorescence microscopy. *A*, 143B-rho^+^ cells expressing TMEM70 cDNA were fixed and immunostained with antibodies against Mitofilin/ Mic60 (upper panel), or TMEM70 (lower panel). Confocal (left) and STED super-resolution (right) images are shown with insets magnified on the right panels. *B*, 143B-rho^+^ cells expressing TMEM70-FLAG were fixed, immunostained with antibodies against Mitofilin/Mic60 (red) and FLAG (green), counter-stained with DAPI (blue), and subjected to expansion microscopy. A representative image is shown (left panel) with an inset magnified on the right panel in separate channels (upper right) and overlay (lower right). *C*, cultured human skin fibroblasts were fixed, immunostained with antibodies against TMEM70 (green), and subjected to expansion microscopy. A representative image is shown on the left and the inset magnified on the top right corner. *D*, scheme representing the relative localization of Mic60 (in red) and TMEM70 (in green) in the inner membrane sub-compartments. OM: outer membrane; IBM: inner boundary membrane; ICM: inner cristae membrane. Scale bars: 5 μm full size images, 500 nm zoomed images (D, 1mm).

**Figure 3.**
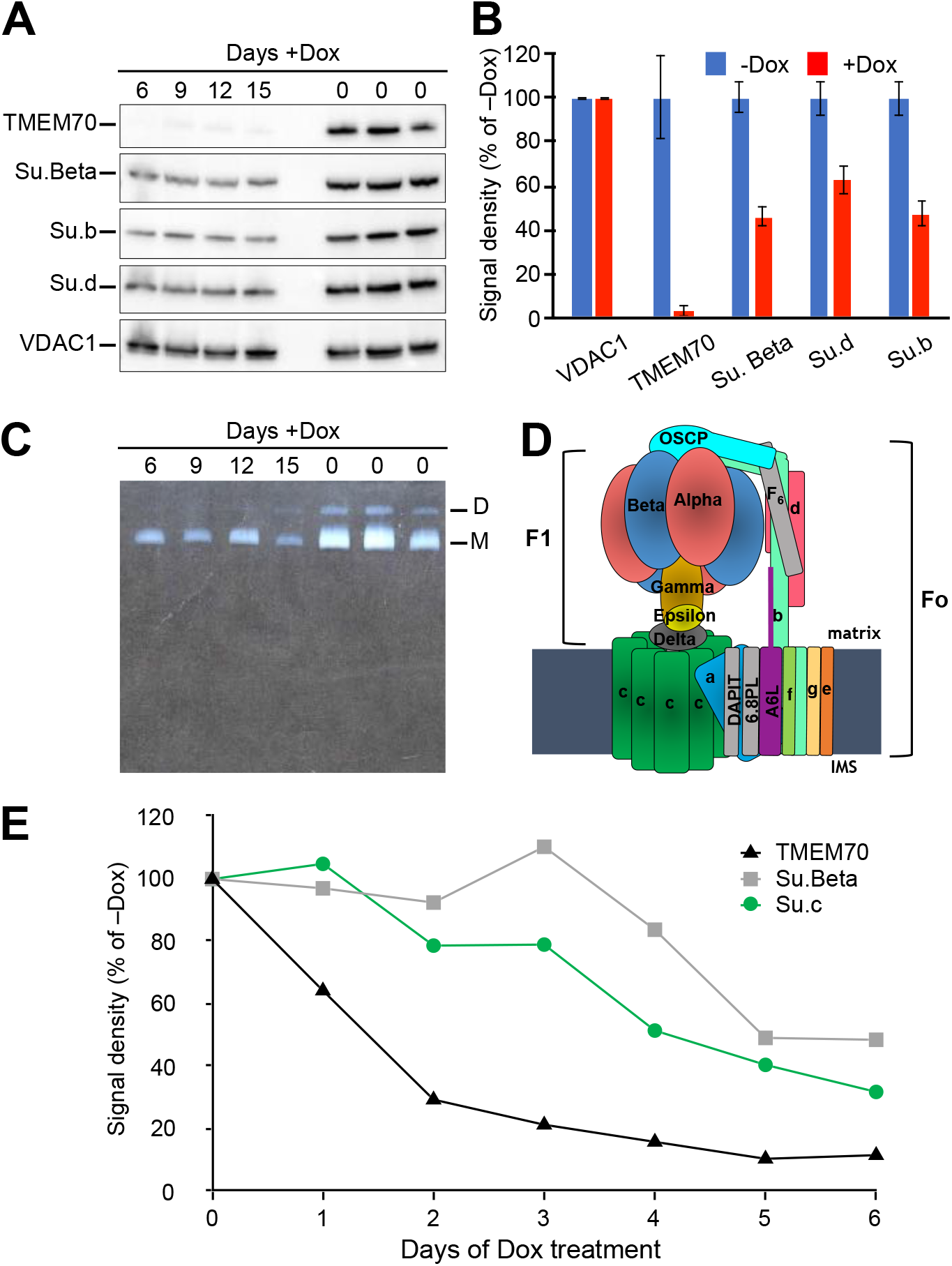
Inducible knockdown of TMEM70. HEK293T cells were transduced with constructs allowing the doxycycline-inducible expression of a mix of two shRNAs targeted to TMEM70 mRNA. *A-C*, cells were cultured in the presence of doxycycline for indicated time periods or left without the drug during equal time and harvested. Mitochondria were isolated from the cell pellets and analyzed as follows. *A*, mitochondria were lysed, and proteins were separated by SDS-PAGE and analyzed by western blot. *B*, quantitation of signal intensities from A. Signals were normalized to VDAC1 used as a loading control. For each protein tested, an average of normalized signals in the presence of doxycycline at the different time points is shown as a percentage of the average signal obtained in the absence of doxycycline; +/− Standard Deviation. *C*, digitonin extracts from the isolated mitochondria were subjected to BN-PAGE and ATPase activity was revealed in-gel (white precipitate). The position of ATP synthase monomers (M) and dimers (D) are indicated to the right. *D*, scheme of the human ATP synthase structure. The general structure of the complex is represented on the scheme with the different types of subunits colored and named. *E*, cells were cultured in the presence of doxycycline for the indicated time periods and harvested. Total protein extracts were separated by SDS-PAGE and analyzed by western blot. Signal densities were quantified and expressed as percentage of signal obtained in the absence of doxycycline.

To better discriminate between the two inner membrane domains, we developed an expansion microscopy approach. By opposition with STED that aims to increase optical resolution of the microscope, the rationale of expansion microscopy is rather to increase the size of the sample and image it with a standard wide-field microscope (46, 47). In this way, the samples were expanded about four-fold in all three dimensions. To enable the parallel localization of TMEM70 and MIC60, we used cells stably expressing endogenous and FLAG-tagged TMEM70 (Fig. 2B). Expansion microscopy confirmed the localization of MIC60 to punctate MICOS structures at the mitochondrial periphery. TMEM70-FLAG did not co-localize to MIC60-positive puncta, but distributed to structures entirely spanning the mitochondrial width as expected for mitochondrial cristae. Interestingly, the TMEM70-FLAG-labelled protrusions often aligned with MIC60, as expected from the localization of MICOS at the junctions between cristae and boundary membranes (See Fig. 2D for a schematic representation). Finally, we localized endogenous TMEM70 with our TMEM70-specific antibodies. The signal to noise was lower, but the images depicted the cristae-like distribution observed for TMEM70-FLAG and no significant signal was observed in the inner boundary membrane (Fig. 2C).

These observations show that TMEM70 is mostly located within the inner cristae membrane (see Fig. 2D), suggesting that the steps of ATP synthase biogenesis mediated by TMEM70 occur within the cristae domains of the inner mitochondrial membrane.

### The knockdown of TMEM70 leads to reduced amounts of functional ATP synthase

To investigate the molecular role of TMEM70 in the assembly pathway of the ATP synthase, we used a knock-down strategy in which TMEM70 was silenced using doxycycline-inducible shRNAs in a stably transduced HEK293T-based cell line. TMEM70 became virtually undetectable already after 6 days of shRNA expression and the levels of several subunits of the ATP synthase tested (see Fig. 3D for a schematic view of the ATP synthase) were stably reduced to 40-60% (Fig. 3AB). Consistently, probing ATPase activity in Blue Native gels revealed lower levels of assembled and functional ATP synthase in the TMEM70 knock-down cells (Fig. 3C).

Next, we investigated the kinetics of TMEM70 disappearance, and its consequences, within the first days of doxycycline treatment (Fig. 3E). The levels of TMEM70 decreased rapidly to 30% after 2 days and to less than 10% after 5 days. By contrast, the levels of Su.Beta and Su.c (that was recently reported to interact with TMEM70) decreased with a slower kinetics. The levels of Su.c started to decrease after 2 days of treatment and declined continuously to less than 40% after 6 days. The levels of su.Beta remained stable during the first 3 days, decreased to 50% between days 3 and 5 and remained stable after day 5 (Fig. 3E).

These results suggest that TMEM70 levels must decrease below some threshold level (~30% of wild-type levels) before the abundance of Su.c and of the ATP synthase become significantly diminished. We thus infer that, while TMEM70 is not absolutely mandatory for the ATP synthase assembly/stability process, it is critical to sustain life-compatible levels of ATP synthase and rates of ATP synthesis.

### The absence of TMEM70 specifically impacts the ATP synthase with a marked decrease in subunit c levels

To further characterize the consequences of the loss of TMEM70, we generated by CRISPR/Cas9 TMEM70 knock-out cells from the HEK293T cell line using sgRNAs targeting either exon1 or exon2 of TMEM70’s gene. Six clones with expected alterations at the TMEM70 genomic locus were retained for further analyses. They all showed undetectable levels of TMEM70, unaffected levels of control proteins (CORE2 subunit of Complex III and the TOM20 subunit of the translocase of the outer mitochondrial membrane translocase), and marked reductions in the levels of several ATP synthase subunits (Fig. 4A).

**Figure 4.**
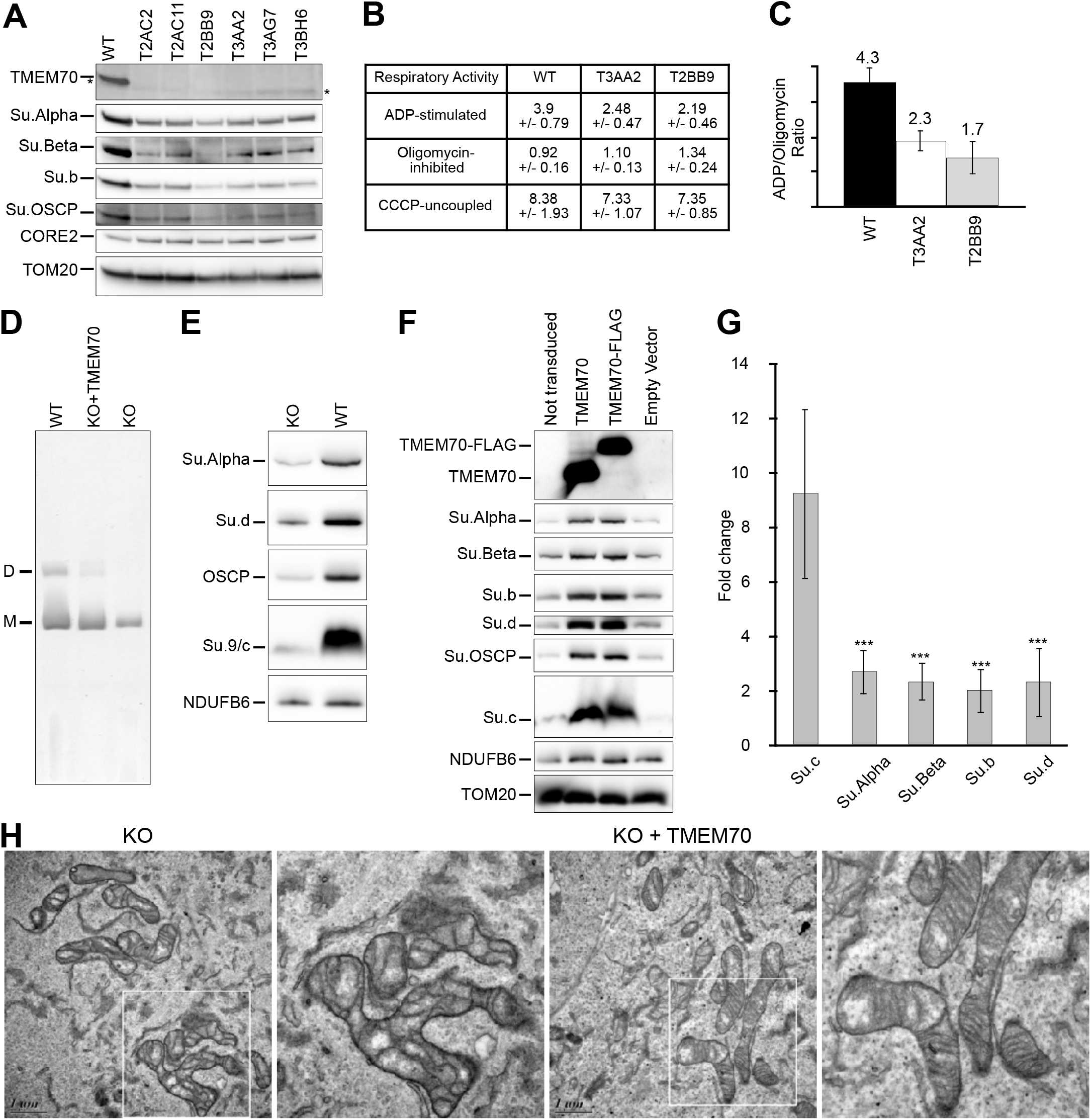
Characterization of TMEM70-knockout cells generated from HEK293T. *A*, isolated mitochondria from different TMEM70-KO clones and HEK293T as a control were lysed and proteins were separated by SDS-PAGE and analyzed by western blot. The faint signal indicated by * just below TMEM70 band is an unspecific cross-reaction of anti-TMEM70 serum. *B*, TMEM70-KO (T3AA2; T2BB9) and HEK293T permeabilized cells were subjected to oxygen consumption measurements after addition of ADP, Oligomycin, and CCCP. The results shown, expressed in nmolO_2_/min/10^6^cells, are average +/− SD of 6 to 9 measurements from 3 to 4 biological replicates, respectively. *C*, the ratio of ADP-stimulated rate over Oligomycin-inhibited rate was calculated for each measurement and the average ratios +/− SD for each cell line are represented in the bar graph with corresponding numerical values indicated above each bar. *D*, digitonin extracts from Isolated mitochondria of wild-type (HEK293T), T3AA2 TMEM70 knock-out (KO), and the same KO cell line expressing exogenous TMEM70 (KO+TMEM70) were subjected to BN-PAGE and ATPase activity was revealed in-gel. ATP synthase monomers (M) and dimers (D) are detected in these conditions. *E*, isolated mitochondria from HEK293T (WT) and TMEM70 knock-out (KO) cell lines were lysed and proteins were separated by SDS-PAGE and analyzed by western blot. *F*, total protein extracts from KO cell lines transduced with the indicated constructs were separated by SDS-PAGE and analyzed by western blot. *G*, quantitation of relative signal intensities of several ATP synthase subunits from E, F, and additional samples for each condition. The results are expressed as the fold change in levels of each subunit when TMEM70 is expressed as compared to KO. Data are mean +/− SD; *n*=6-16. \*\*\**P*<0,001 ANOVA. *H*, TMEM70 KO cells expressing TMEM70 cDNA (KO + TMEM70) or not (KO) were fixed, sliced and imaged by transmission electron microscopy. Representative images are shown in the upper panels and the insets are magnified in the lower panels.

Two clones, T3AA2 and T2BB9, were subjected to oxygen consumption assays in digitonin-permeabilized cells (Fig. 4B). Three respiratory states were assessed: State 3 or phosphorylating state, in the presence of a saturating amount of ADP; Oligomycin state where ATP synthase is fully inhibited and the respiratory rate is minimal (compensates only the membrane passive permeability to protons); and the CCCP uncoupled state where respiratory rate is maximal in the absence of upstream kinetic control. State 3 respiration was decreased in both knock-out cell lines to a comparable extent when compared to wild-type HEK293T cells, whereas no significant difference between the mutant and control cells was observed in the oligomycin-inhibited and uncoupled states. Noteworthy, knock-out cells displayed an important 50-60% decrease in the ratio of state III over oligomycin state rates indicating a specific ATP synthase deficiency (Fig. 4C). As the characterized TMEM70 knock-out clones gave similar results, only one (T3AA2, hereafter designated as KO) was retained for all further experiments described below.

We first investigated the assembly status and abundance of the ATP synthase in the KO cell line by probing ATPase activity in Blue Native gels, and the steady state levels of individual ATP synthase subunits by western blot analysis of denaturing gels. Consistent with the respiration assays, these analyses revealed reduced amounts of assembled ATP synthase (Fig. 4D) and of individual subunits in the KO cells (Fig. 4EG) that were reversed to normal levels upon transduction with untagged or FLAG-tagged TMEM70 (Fig. 4FG). Interestingly, among the tested ATP synthase subunits, the content in Su.c was the most affected by the absence of TMEM70 (Su.c 9-10-fold reduced levels versus 2-3-fold reduced levels of other subunits; see Fig. 4G), which is a unique feature among ATP synthase assembly-defective mutants (see introduction). In accordance with the bioenergetic demise and the role of the ATP synthase in shaping/maintaining cristae, mitochondrial ultrastructure was altered in the KO cells with less cristae and sometimes aberrant shape, and these defects were reversed upon restoration of TMEM70 expression (Fig. 4H).

### Analysis of patient fibroblasts lacking the C-terminal hydrophilic domain of TMEM70

To probe the importance of the hydrophilic C-terminal domain of TMEM70, we analyzed cultured skin fibroblasts from a patient with a homozygous frame-shift mutation (c.497_498del) resulting in truncated TMEM70 after its second transmembrane segment (p.Tyr166Cysfs*7, Fig. 5A). No signal for TMEM70 could be detected by western blot using the rabbit anti-peptides serum described above, indicating that the truncated protein was unstable and degraded (Fig. 5B and S3). Several ATP synthase subunits were in strongly decreased amounts relative to control cells, such as the Su.c (Fig. 5BC) and mitochondrial ultrastructure was altered with much less numerous and irregular arrangement of cristae and a matrix less dense to electrons (Fig. 5D). All these alterations were efficiently suppressed upon transduction of the patient’s fibroblasts with TMEM70 lentiviral construct.

**Figure 5.**
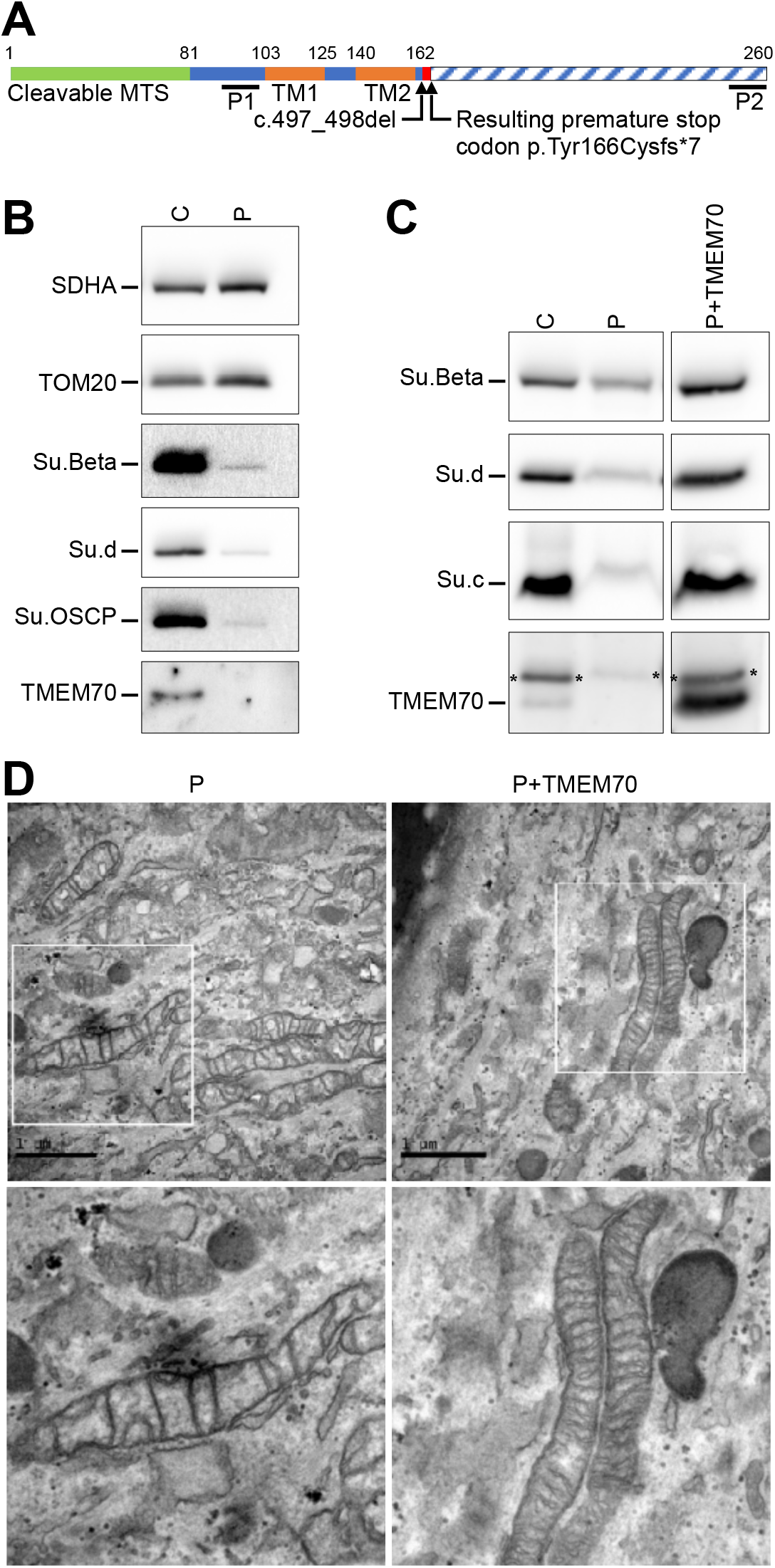
Characterization of patient skin fibroblasts bearing the c.497_498del mutation in TMEM70. *A*, scheme representing the truncated version of TMEM70 resulting from the *c.497_498del* mutation leading to premature stop codon p.Tyr166Cysfs*7. MTS: Mitochondrial Targeting Sequence. TM: Transmembrane segment. *B*, isolated mitochondria from cultured fibroblasts of the patient (P) and a control individual (C) were lysed and proteins were separated by SDS-PAGE and analyzed by western blot. *C*, total protein extracts from control fibroblasts (C) and patient fibroblasts transduced with the empty vector (P) or expressing exogenousTMEM70 (P+TMEM70)were separated by SDS-PAGE and analyzed by western blot. Contaminant signal surrounded by * represents residual signal from Su.d in the subsequent decoration of TMEM70. *D*, patient fibroblasts expressing TMEM70 (P+TMEM70) or GFP as a control (P) were fixed, sliced and imaged by TEM. Representative images are shown.

These results further corroborate the data obtained with KO cells and provide evidence that a compromised ability to assemble the ATP synthase was responsible for the disease of the patient with the c.497-498del mutation in TMEM70. They also underscore the importance of the C-terminal domain of TMEM70 for assembling the ATP synthase.

### TMEM70 specifically protects subunit c from degradation regardless of other ATP synthase subunits and of the energetic status of mitochondria

To gain further insights into the role of TMEM70 in ATP synthase assembly, we investigated the levels of ATP synthase subunits in cells endogenously expressing TMEM70 but devoid of mtDNA (Rho°) and thus missing core components of the respiratory chain and the two mtDNA-encoded Fo subunits Su.a and Su.A6L. We found that, similarly to TMEM70 KO cells, the levels of Su.c were significantly reduced compared to other ATP synthase subunits in the Rho° cells (Fig. 6A). This observation raised the possibly that the lack of Su.c in TMEM70 KO cells was not primarily caused by the absence of TMEM70. However, we found that, similarly to Su.c, the levels of TMEM70 were also significantly reduced in Rho° cells (Fig. 6A). Interestingly, when the Rho° cells were transduced with our TMEM70 construct, normal steady state levels of Su.c were recovered (Fig. 6B). The poor accumulation of Su.c in cells lacking mtDNA is thus due to the decreased abundance of TMEM70 in these cells highlighting a reciprocal stabilization of Su.c and TMEM70.

**Figure 6.**
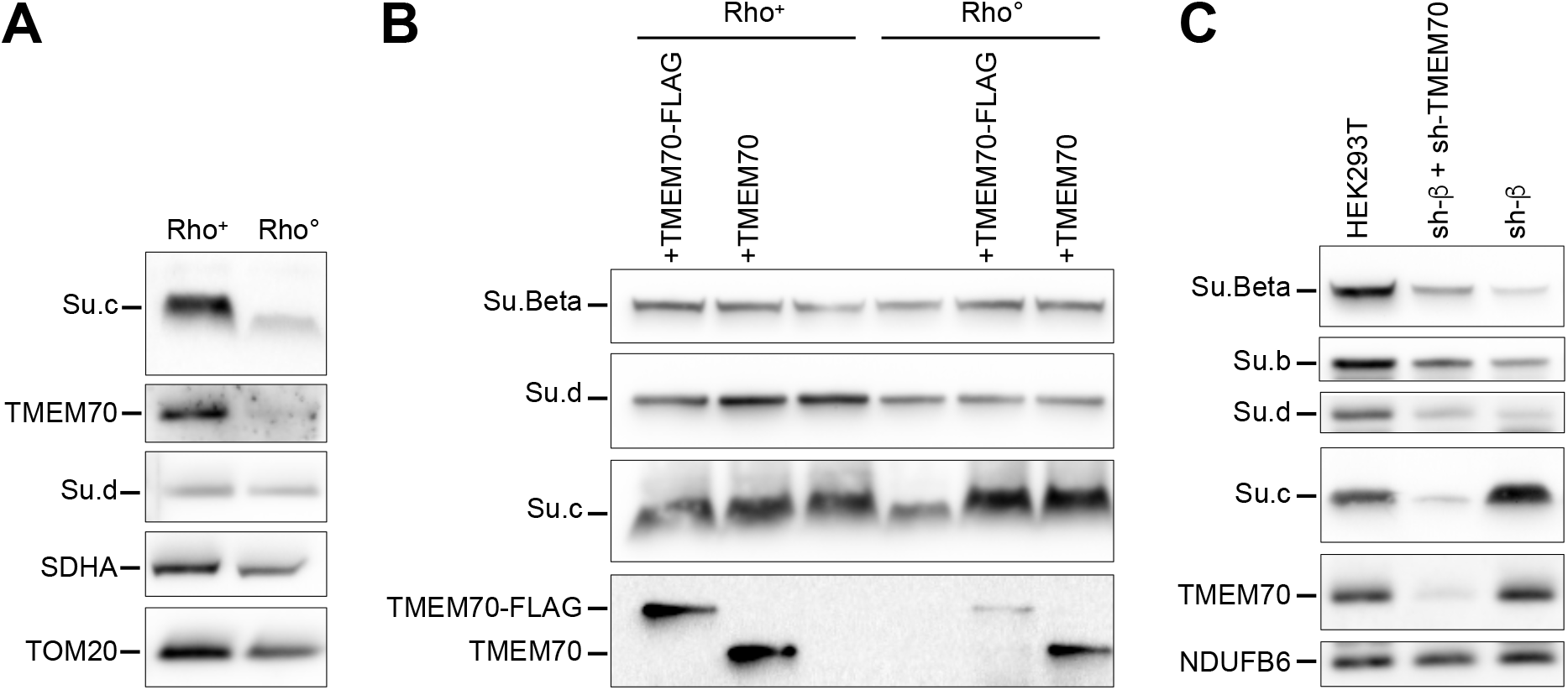
TMEM70 targets specifically the subunit c of the ATP synthase. *A*, isolated mitochondria from 143B cell line containing (Rho^+^) or not (Rho°) mitochondrial DNA were lysed and proteins were separated by SDS-PAGE and analyzed by western blot. *B*, total protein extracts from 143B cell line containing (Rho^+^) or not (Rho°) mitochondrial DNA expressing the indicated constructs were separated by SDS-PAGE and analyzed by western blot. *C*, HEK293T cells were transduced with shRNA constructs to knock-down TMEM70 (sh-TMEM70) and/or ATP5B (sh-β). Non-transduced HEK293T were used as control. Total protein extracts were prepared, separated by SDS-PAGE and analyzed by western blot.

Next, we aimed to know whether the F1 subcomplex, which is entirely encoded by nuclear genes, also influences the stability of TMEM70. To this end, we knocked-down Su.Beta in rho^+^ cells (Fig. 6C). While the Fo subunits b and d paralleled the depletion in F1 Su.Beta (indicating a major block in the assembly of the ATP synthase holoenzyme), the levels of Su.c, TMEM70, and a component of the respiratory complex I (NDUFB6) were not affected. The amounts of Su.c were reduced only when both TMEM70 and Su.Beta were knocked-down concomitantly (Fig. 6C).

Taken together, these experiments revealed that TMEM70 is required to maintain the levels and stability of Su. c regardless of other ATP synthase subunits and of the energetic and genetic status of mitochondria. They further suggest that TMEM70 specifically protects subunit c from degradation indicating that the sharp reduction in fully assembled ATP synthase in cells lacking TMEM70 results from a higher susceptibility to degradation of Su.c.

### TMEM70 forms homo-oligomers interacting transiently with subunit c before its incorporation into the ATP synthase

To gain further insights into the links between TMEM70 and Su.c, we isolated mitochondria from TMEM70 knock-out cells expressing untagged or FLAG-tagged TMEM70, solubilized these mitochondria using digitonin (a mild detergent that preserves protein-protein interactions), and performed co-immunoprecipitation tests using TMEM70-FLAG as a bait. TMEM70-FLAG was almost entirely immunoprecipitated from the digitonin extracts using beads covalently bound to anti-FLAG monoclonal antibodies (Fig. 7A). Su.c, but none of the other tested ATP synthase subunits, repeatedly co-immunoprecipitated with TMEM70-FLAG although in very small amounts. Noteworthy, in control experiments with non-tagged TMEM70, neither TMEM70 nor Su.c was pulled down. Since most Su.c was in the flow through, as other ATP synthase subunits (Fig. 7A: FT), we infer that TMEM70 transiently interacts with the low levels of Su.c not yet incorporated into the ATP synthase. Remarkably, similar results were also obtained with cells devoid of mtDNA (Fig. S4), indicating that mtDNA-encoded proteins, including Fo transmembrane subunits a and A6L, do not interact with TMEM70 and are not required for co-immunoprecipitation of TMEM70 and Su.c.

**Figure 7.**
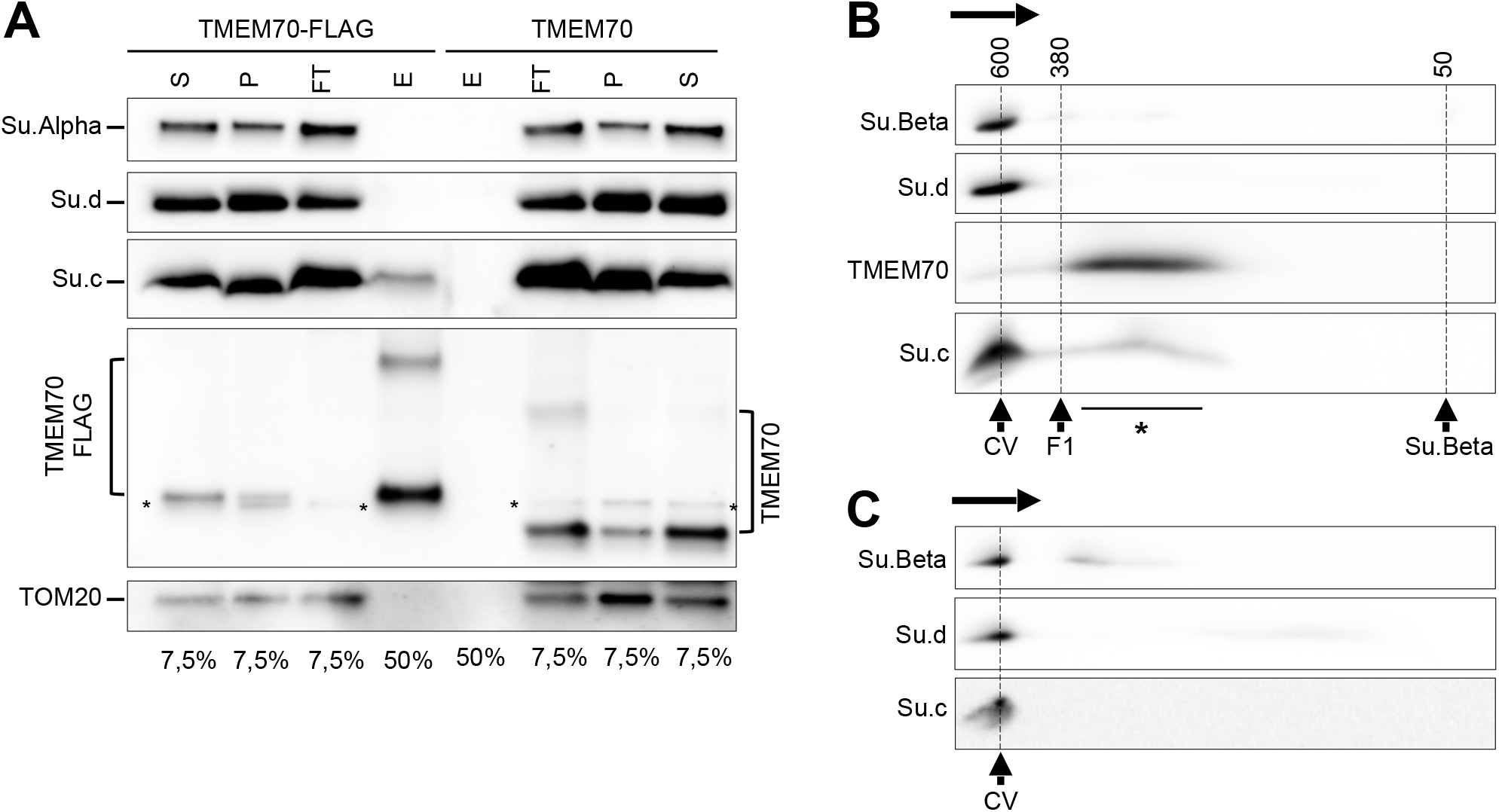
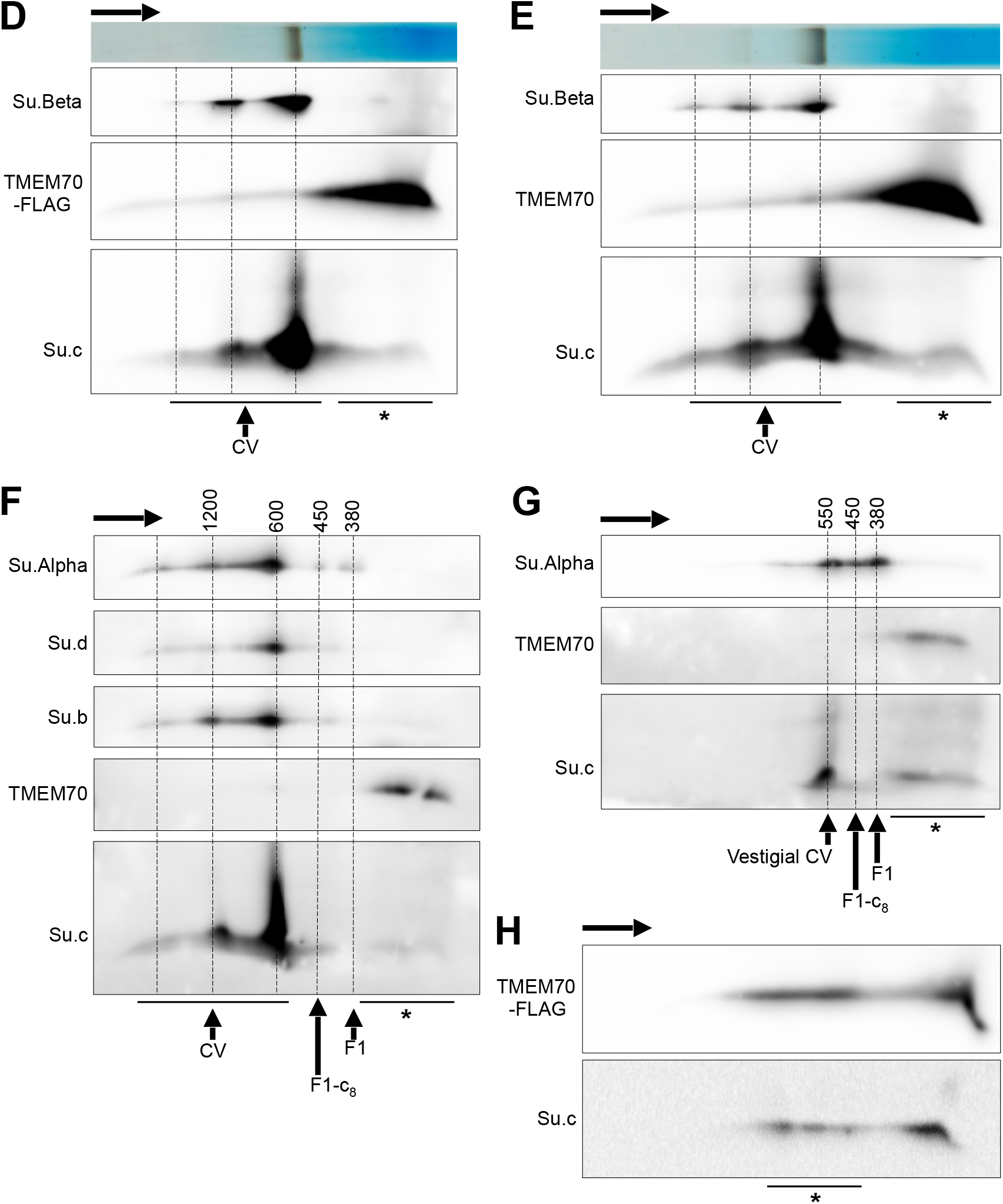
Physical interaction between TMEM70 and the subunit c of the ATP synthase. *A*, immunoprecipitation of TMEM70-FLAG. Mitochondria isolated from TMEM70-KO cells overexpressing TMEM70-FLAG, or TMEM70 as a control, were used in immunoprecipitation assays using anti-FLAG M2 magnetic beads. Digitonin extractions were performed in order to preserve protein-protein interactions. Extraction supernatants (S) represent the input material incubated with the beads while insoluble pellet (P) were set apart for analysis. The unbound material come with the flow-through (FT) while bound proteins are retained on the beads and further eluted (E) natively by competition with free 3xFLAG peptide. Portions (indicated at the bottom of the panel) of each fraction were analyzed by SDS-PAGE and western blot. The signals surrounded by * represent residual signal from Su.d in the subsequent decoration of TMEM70. *B-H*, mitochondria were isolated for each cell line and subjected to digitonin extraction. The extracts were analyzed by BN-PAGE in a first dimension. The horizontal arrow above each panel indicated the migration direction. Some lanes were used for in-gel ATPase activity assay where shown. All other lanes of the BN-PAGE were cut and analyzed by SDS-PAGE in a second dimension. Proteins were transferred to nitrocellulose and analyzed by western blot. Position of fully (CV) or partially (Vestigial CV) assembled ATP synthase, F1-c_8_ subcomplex, F1-ATPase, and free Su.Beta, or TMEM70-containing complexes (*), are indicated below each panel with approximate calculated molecular weight in KDa above panels when applicable. *BC*, analysis of original HEK293T (B) and TMEM70 KO (C). The high molecular weight part of the 4-16% BN-PAGE lanes was cut in order to fit lanes till bottom in the SDS-PAGE 2D well. *D-G*, the low molecular weight part of the 3-12% BN-PAGE lanes was cut in order to fit lanes till top in the SDS-PAGE 2D well. *DE*, analysis of TMEM70 KO expressing TMEM70-FLAG (D) or TMEM70 (E). These samples were loaded twice on the BN-PAGE. One lane was used for 2D western blot analysis and the second was used to reveal in-gel ATPase activity that is shown at the top of each panel. *FG*, analysis of 143B cell line containing (Rho^+^, F) or not (Rho°, G) mitochondrial DNA. *H*, native eluates from the immunoprecipitation of TMEM70-FLAG described in A were analyzed by two-dimensional electrophoresis and western blot, as described above for mitochondrial extracts. TMEM70-Su.c complexes display different sizes (*), probably reflecting increasing incorporation of Su.c molecules.

An additional, weaker, immunological signal for TMEM70-FLAG was observed in our denaturing gels with approximately a doubled size suggesting that TMEM70 can form dimers, as previously reported (39, 41), that are relatively resistant to detergents (Fig. 7A). It is to be noted that untagged TMEM70 found in the flow-through of control experiments shows a similar additional signal (considering the size of the tag), ruling out any artifact resulting from the tagged version. This result suggests that higher molecular weight complexes of TMEM70 interact with Su.c.

To further define the interactions between these proteins, we separated digitonin-solubilized protein complexes by two-dimensional electrophoresis (BlueNative/BN-PAGE followed by denaturing/SDS-PAGE) and analyzed them by western blot. In extracts from HEK293T control cells, as expected, ATP synthase subunits of the F1, proton channel, and peripheral stalk (Su.Beta, Su.c, Su.d, respectively) co-migrated at the position of fully assembled complex V (Fig. 7B). Note that ATP synthase monomers and oligomers are not well resolved in this 4_16% polyacrylamide native gel. All TMEM70 signal was in smaller size complexes of grossly 300-400 kDa (as compared to the calculated molecular weights of the F1 particle and free su.Beta detected with Su.Beta antibodies). This provides evidence for endogenous TMEM70 oligomers much beyond dimers. In addition, these results further demonstrate that TMEM70 does not associate with fully assembled ATP synthase, in agreement with Fig. 7A.

Interestingly, in addition to the strong Su.c signal co-migrating with the ATP synthase, an additional Su.c signal of weaker intensity co-migrated with the protein complexes responding to TMEM70 antibodies. As expected, the Su.c signal exclusively co-migrated with Complex V in extracts from the TMEM70 KO clone (T3AA2) (Fig. 7C) albeit in reduced amounts as compared to control cells, consistent with the reduced amounts of complex V in the KO cells (Fig. 7BC). This confirmed the results of our pull-down assays and provide the first direct experimental evidence that only Su.c not yet incorporated into the ATP synthase interacts with TMEM70.

Similar results were obtained with KO cell lines expressing either a FLAG-tagged version or an untagged version of TMEM70 (Fig. 7DE). Of note, in the experiments shown in Fig. 7DE, 3-12% polyacrylamide native gels were used allowing to resolve ATP synthase monomers and oligomers. Interestingly, longer western blot exposures (Fig. S5) revealed an additional weaker signal corresponding to TMEM70 dimers similar to that observed in our immunoprecipitation assays.

To deepen our study, we then analyzed samples from the rho^+^ cell line 143B and a Rho° derivative thereof both endogenously expressing TMEM70. With the rho^+^ cells (Fig. 7F), the results were similar to those obtained with wild-type HEK293T cells (Fig. 7B) or with TMEM70 KO cells expressing exogenous TMEM70-FLAG or untagged TMEM70 (Fig. 7DE). Consistent with previous studies (16, 27, 32), partial assemblies that are barely detectable in rho^+^ cells accumulated in rho° cells (F1 and F1-c subcomplexes), together with a so-called ‘vestigial’ complex containing all subunits except for the two mtDNA-encoded ones (a and A6L), DAPIT, and 6.8PL (Fig. 7FG). Interestingly the TMEM70-containing complexes and their co-migration with unassembled Su.c were still observed in these cells (Fig. 7G). These data further show in a more direct manner that none of the proteins encoded by mtDNA, notably Fo Su.a and Su.A6L, is required for the interactions between TMEM70 and Su.c.

Furthermore, 2D analysis of immunoprecipitated TMEM70-FLAG revealed that high molecular weight complexes resisting precipitation and elution depict several discrete spots of Su.c with increasing molecular weights (Fig. 7H). The existence of complexes with different TMEM70-Su.c stoichiometries points to the gradual incorporation of c-subunits to TMEM70 oligomers and hence suggests a role for TMEM70 in the assembly of the c-ring (a homo-oligomeric ring of 8 Su.c molecules).

## Discussion

Although the involvement of TMEM70 in mammalian ATP synthase biogenesis is unquestioned (34, 36, 40, 48–50), its precise molecular function remained largely unknown. We herein provide evidence that TMEM70 localization is restricted to cristae where it assembles in large oligomers that interact specifically with Su.c and provide a scaffold for the assembly of the c-ring before its incorporation into functional F1Fo ATP synthase.

We report that the lack of TMEM70 leads to a specific deficiency of the ATP synthase via a drastic drop in Su.c levels (Fig. 3–5), further corroborating a recent study that has revealed a role of TMEM70 in the biogenesis of Su.c (37). We also show that the loss of mitochondrial DNA (rho^0^) and hence of mtDNA-encoded subunits, known to hamper the assembly of ATP synthase, leads to a significant decrease in the levels of nuclear-encoded Su.c and, interestingly, of TMEM70 (Fig. 6AB). Noteworthy, expression of TMEM70 in rho^0^ cells restored the levels of Su.c without affecting those of other subunits. We additionally show that the Su.c levels were not impacted in rho^+^ cells unable to express the F1-Beta protein (Fig. 6C), similarly to cells in which one of the central stalk subunits of the F1 was silenced (24, 26) and in agreement with results obtained with F1-Beta−/− cells (37). Moreover, Su.c levels were drastically reduced only when F1-Beta and TMEM70 were concomitantly knocked-down (Fig. 6C). We infer that none of the subunits of respiratory chain complexes and ATP synthase encoded by the mtDNA, nor F1 subunits or even the energetic status of mitochondria, directly influence the levels of Su.c. Instead, the levels of Su.c appear to directly and specifically correlate with those of TMEM70 within mitochondria.

Recently, the existence of physical interactions between TMEM70 and Su.c was revealed by pull down assays using a FLAG-version of su.c (37). In these assays the majority of overexpressed FLAG-tagged Su.c was immunoprecipitated, together with substantial proportions of endogenous Su.c and TMEM70. Of note, this experimental design did not allow to establish the fraction of endogenous Su.c interacting with TMEM70. In contrast, our immunoprecipitation assays using TMEM70 as a bait showed that only a minor amount of the pool of endogenous Su.c was pulled down, and that none of the other tested subunits of ATP synthase co-precipitated (Fig. 7A). Furthermore, in native gels, TMEM70 formed 300-400 kDa protein complexes that co-migrated with a fraction of Su.c not incorporated into ATP synthase (Fig. 7B-H). The present study therefore provides the first and robust evidence that endogenous TMEM70 forms large oligomers, further corroborated by results obtained with FLAG-tagged TMEM70 before or after immunoprecipitation (Fig. 7). Furthermore, the present study provides first evidence that TMEM70 interact with Su.c as an oligomeric structure and provides direct evidence that it only makes contacts with unassembled Su.c prior to its association with its protein partners within the ATP synthase. Our observations of 300-400 kDa TMEM70 complexes that interact with free Su.c contrast with previous studies reporting interaction of TMEM70 with partial ATP synthase assemblies (36) or the assembly of TMEM70 into small complexes likely representing dimers (37, 39, 41).

Consistent with previous reports (39–41), our biochemical studies localize endogenous TMEM70 in the mitochondrial inner membrane with the loop connecting its two transmembrane segments in the intermembrane space and the C-terminal hydrophilic domain on the other side of the membrane in the matrix (Fig. 1). Interestingly, not only the hydrophilic C-terminal domain is evolutionary well conserved but the primary sequence of the transmembrane segments is particularly conserved too (Fig. S6), indicating that they do not simply represent membrane anchors. Possibly the transmembrane segments of TMEM70 are crucial for function and for specific interactions with Su.c. Another interesting feature is the ability of TMEM70 molecules to form stable dimers (39, 41) that resist dissociation even under denaturing conditions (Fig. 7A and S5).

Based on the molecular weight of TMEM70-containing complexes (Fig. 7), the known stoichiometry of eight Su.c within the c-ring, and the co-migration of the TMEM70 complexes with Su.c, we propose that TMEM70 is arranged in an octameric structure of eight TMEM70 dimers of 40 kDa (320 kDa) that could interact with up to eight subunits c (380 kDa) (Fig. 8A). We therefore conclude that TMEM70 provides a scaffold easing the formation of the c-ring that is released after completion and incorporation of the c-ring into the ATP synthase to accommodate newly imported Su.c. In support to this hypothesis, the 2D analysis of our immunoprecipitates revealed a series of discrete TMEM70-Su.c complexes with apparent enrichment in Su.c content along with increasing molecular weight (Fig. 7H).

**Figure 8.**
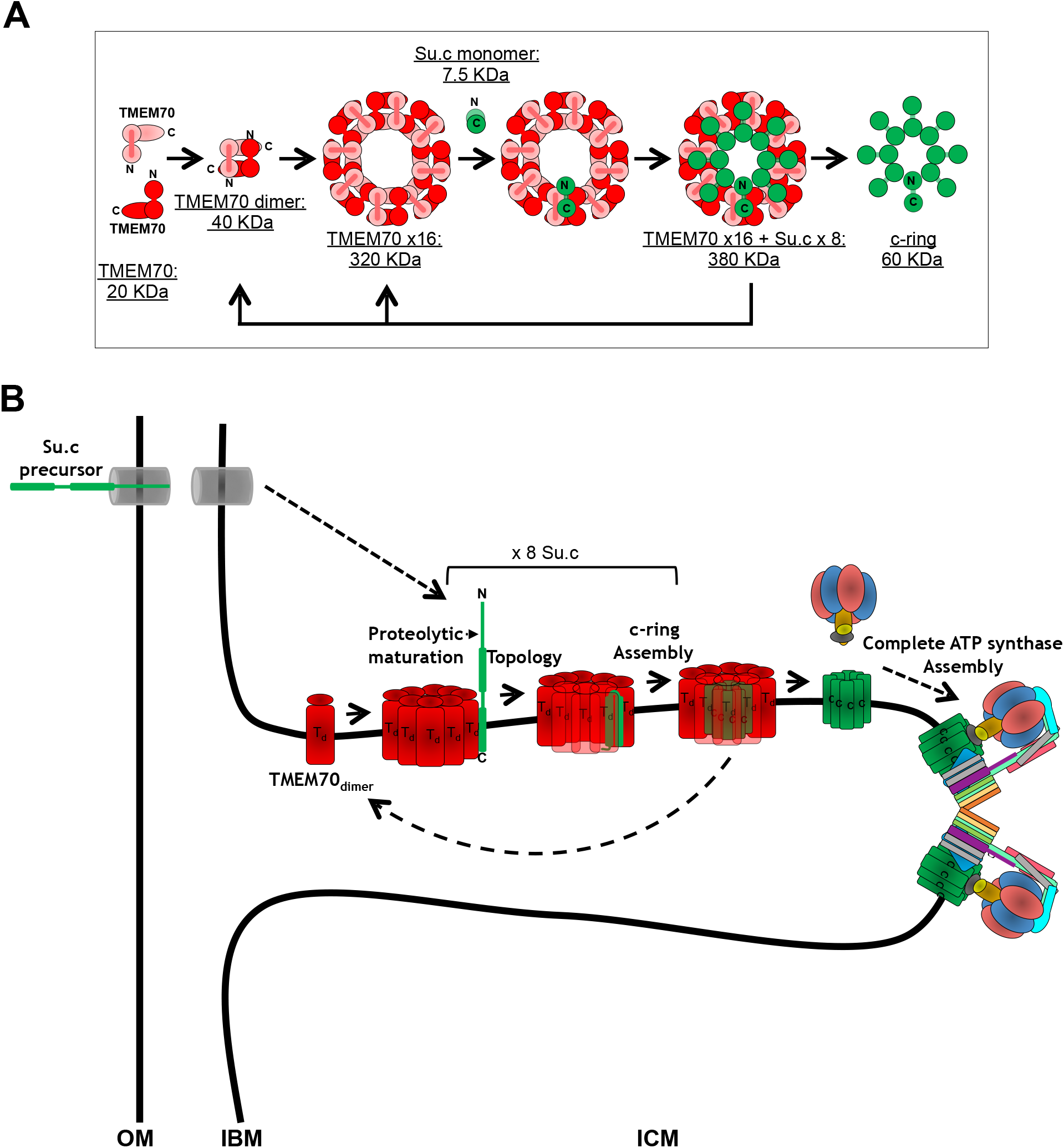
Model for the molecular action of TMEM70 towards the subunit c of the ATP synthase. *A*, molecular model for the TMEM70-dependent assembly of the c-ring. TMEM70 forms dimers that arrange in eight-copy structure in which each dimer can accommodate one Su.c in order to form the c-ring. After the c-ring is released, TMEM70 structural ring or disassembled dimers are involved in a new c-ring formation as indicated by the arrows. Approximate calculated molecular weights for each structure is indicated on the figure. *B*, the biogenesis of the ATP synthase is represented with a special focus on the c-ring. The Su.c precursor (in green) is imported into mitochondria through the protein translocases (grey cylinders) across the outer membrane (OM) and the inner boundary membrane (IBM). The Su.c precursor gets processed by the Matrix Processing Peptidase (Proteolytic maturation). TMEM70 (in red) forms dimers (TMEM70_dimer_ or T_d_) in cristae that interact with Su.c within higher order oligomers in order to form the c-ring. TMEM70 complexes release directly or disassemble into dimers to liberate the c-ring and take in charge new Su.c. The c-ring next assembles with premade F1 subcomplex and the additional subunits are added to form ATP synthase monomers, dimers, and oligomers.

The OXPHOS complexes were shown, notably in mammals, to accumulate in cristae with a differential enrichment of respiratory chain complexes within the laminar part of cristae and of ATP synthase at the edge of cristae (6, 7, 9, 42, 43). What are the mechanisms and operating forces underlying such a compartmentalization is a challenging question.

The expansion microscopy (46, 47) here applied to mitochondria provides a possible clue to this membranous partition of protein complexes. Indeed, these experiments revealed that TMEM70 localizes in cristae domains of the inner membrane (Fig. 2). These observations strongly suggest that formation of the c-ring, an early step in the assembly process of ATP synthase, takes place within cristae as depicted in Fig. 8B rather than close to the MICOS-dependent contact sites between outer and inner membranes.

Interestingly, a recent study (14) in *S. cerevisiae* (where the subunit c of the ATP synthase is encoded by the mtDNA) showed by immunogold electron microscopy that three factors specifically involved in the translation and assembly of mtDNA-encoded subunits of the ATP synthase were enriched in cristae, by contrast with factors involved in translation and assembly of complexes III and IV.

In mammals, where none of the yeast factors investigated are evolutionary conserved, and where Su.c is nuclear-encoded with additional constraints for its biogenesis (see below), we provide the first evidence for in situ assembly of the ATP synthase within cristae. These topographic differences in the biogenesis of OXPHOS complexes may contribute to the preferential accumulation of Complex V at the edge of cristae, which is presumably important for shaping the mitochondrial inner membrane (6, 7, 9). Consistently, mitochondrial ultrastructure was found perturbed in our TMEM70 KO cells and in cells from a patient carrying a premature stop codon after TM2 that destabilizes TMEM70 (Fig. 4 and 5), similarly to tissues from a mouse lacking this protein (50) and other TMEM70 patients (48, 49, 51).

In yeast, Su.c would insert co-translationally into the membrane with the assistance of the conserved protein translocase Oxa1 (52, 53) and other yeast specific factors (Aep1, Aep2 and Atp25) specifically involved in the synthesis and assembly of Su.c (54–56). These factors are supposed to protect Su.c, which is an extremely hydrophobic protein referred to as a proteolipid, from exposure and non-productive interactions in the aqueous mitochondrial matrix. With the relocation into the nucleus of the Su.c gene in mammals and a few other organisms including filamentous fungi (57–59), the sorting of Su.c to its final destination from the cytosol is more complex, and this has likely required specific adaptive evolution. For instance, evidence was provided that a functional allotopic expression of Su.c from the nucleus in yeast necessitates reducing the hydrophobicity of its first transmembrane segment presumably to avoid its lateral sorting in which case Su.c would be unable to reach a functional topology with its C- and N-termini in the intermembrane space (60).

Therefore, TMEM70 may be regarded as a factor whose emergence was required in mammals to take in charge Su.c after its transport into the organelle to protect it from degradation.

The mitochondrial genome is a remnant of an ancestral prokaryotic genome that has considerably reduced in size during eukaryotic evolution due to a massive transfer of genes to the nucleus. To date, the main function of mitochondrial genomes is the production of a few subunits of the mitochondrial energy-transducing system. Some are always found in the mtDNA of respiring eukaryotes, like the ones encoding the cytochrome b subunit of Complex III and the Cox1 subunit of Complex IV; while others can be found in the nucleus or inside the organelle, like the gene encoding Su.c. The complexity of the functional relocation to the nucleus of this gene in yeast (60) suggests that there must be some compelling reason to have it expressed from nuclear DNA. In support to this hypothesis, it was found that after its natural nuclear relocation in *Podospora anserina*, Su.c production and the efficacy of oxidative phosphorylation became highly regulated along the complex life cycle of this organism from two paralogous genes encoding structurally different subunits c (61, 62).

In humans where Su.c is also expressed from the nucleus, there are three Su.c gene isoforms among which one (P1) shows transcriptional regulations in a cell- and tissue-specific manner, suggesting that, as in filamentous fungi, the Su.c became also a key regulatory target to modulate ATP production in humans. Studies by the group of J. Houstek have provided strong support to this hypothesis (63, 64). The important variations in the content of TMEM70 in mouse tissues reported recently (37), together with the data reported in the present study, suggest that TMEM70 may play a key role in the modulation of ATP synthase biogenesis by controlling the stability of Su.c in mitochondria.

## Experimental Procedures

### Cell lines and culture conditions

HEK293T were used throughout this study. 143B rho^+^ (143B.TK-) is a human osteosarcoma-derived cell line (ATCC CRL 8303). 143B rho° (143B206) is an mtDNA-less derivative of 143B.TK-(65). Both 143B derived cell lines were from Dr. Anne Chomyn (Caltech, USA). Primary human skin fibroblasts from a control individual and from a patient with cardiomyopathy carrying a homozygous frame-shift mutation c.497_498del in the TMEM70 gene were obtained from the Genetic unit of Necker-Enfants malades hospital (Paris, France) and were always kept at early passages. All cell lines were grown in Dulbecco’s modified Eagle’s medium (DMEM; Life Technologies) containing 4.5 g/l glucose, supplemented with 10% fetal bovine serum, 2 mM L-glutamine, 1 mM pyruvate, at 37°C under 5% CO_2_. 50 μg/ml Uridine were added to this culture medium to grow 143B Rho+ and Rho° cell lines.

### Plasmid constructs

TMEM70 cDNA was obtained by PCR amplification from HeLa cDNA preparation. Two fragments were amplified using the primers TMEM70-Fwd and TMEM70-noSfi-Rev or TMEM70-noSfi-Fwd and TMEM70-Rev-Stop in order to introduce a silent mutation (G>A) at position 141 of the coding sequence removing the internal *Sfi*I restriction site naturally present in TMEM70 coding sequence. The two fragments were fused by PCR to obtain a product corresponding to the 783 bp coding sequence of TMEM70 transcript variant 1 that was cloned into pMOS*blue* vector using the Blunt-Ended PCR Cloning Kit (GE Healthcare Life Sciences) to generate the pMOS-TMEM70 plasmid. The insert was subcloned into pKSPS (derived from pBluescriptIIKS(+) by inserting a *Sfi*I and *Pme*I sites in original *Kpn*I and *Sac*I sites, respectively) using the *Xho*I and *Eco*RI sites to generate the pKSPS-TMEM70 plasmid. As to generate a C-terminally FLAG-tagged version, the nucleotide sequence encoding the 3xFLAG epitope (DYKDDDDKDYKDDDDKDYKDDDDK) was obtained by annealing the two primers 3xFLAG-sense and 3xFLAG-antisense and cloned into the *Eco*RI and *Bam*HI sites of the pKSPS vector to generate the pKSPS-FLAG plasmid. TMEM70 cDNA was amplified by PCR from pKSPS-TMEM70 using a different primer at the 3’ end (TMEM70-Rev-NoStop) removing the stop codon and the fragment obtained was cloned into the *Xho*I and *Eco*RI sites of pKSPS-FLAG, in frame with the 3xFLAG tag sequence, to generate pKSPS-TMEM70-FLAG. The TMEM70 and TMEM70-FLAG inserts were then subcloned into the previously described lentiviral vector pCMV-IRES-mitoDsRed (38) using the *Sfi*I and *Pme*I sites to generate the pCMV-TMEM70-IRES-mitoDsRed and pCMV-TMEM70_FLAG-IRES-mitoDsRed plasmids, respectively.

For TMEM70 knockdown studies, target sequences in TMEM70 mRNA were chosen (T1 GGGAAGGATATGTTCGATTCTTAAA and T2 CGAGTCTGATTGGCCTTACATTTCT) and primers were designed to encode shRNAs (shT1-Fwd/shT1-Rev and shT2-Fwd/shT2-Rev). For each construct, complementary primers were hybridized and cloned into the *Bam*HI and *Eco*RI sites of the lentiviral pSIH-H1-copGFP-shRNA vector (System Biosciences) according to the manufacturer’s instructions to generate pSIH-T1 and pSIH-T2 plamsids for constitutive knockdown. In order to achieve doxycycline-inducible knockdown, a different set of primers targeting same regions of TMEM70 mRNA was used (shT1-Fwd-pLV, shT1-Rev-pLV, shT2-Fwd-pLV, shT2-Rev-pLV) to clone shRNA encoding sequences into the *Mlu*I and *Cla*I sites of the lentiviral pLVTHM vector (66) in order to generate pLVTHM-T1 and pLVTHM-T2. pLVTHM and pLV-tTR-KRAB-red were gifts from Didier Trono (Addgene plasmids # 12247 and # 12250).

To perform the Su.Beta knockdown, TRC2-pLKO-puro-shBeta3 (NM_001686.3-422s21c1) and TRC2-pLKO-puro-shBeta4 (NM_001686.3-1488s21c1) lentiviral constructs (Sigma-Aldrich) expressing shRNAs targeting the ATP5B mRNA were used.

To generate TMEM70 Knockout cell lines using the CRISPR-Cas9 technology, four different sgRNAs designed previously for a genome-scale CRISPR-Cas9 knockout (GeCKO) screening (67) were used, two (TMEM70_sgRNA-1 and TMEM70_sgRNA-2) targeted to the first exon and two others (TMEM70_sgRNA-3 and TMEM70_sgRNA-4) targeted to the second exon of the TMEM70 gene. The dual vector GeCKO system was used consisting of plentiCas9-Blast and lentiGuide-Puro (68). For each construct, a pair of primers was hybridized and cloned into the BsmBI site of pLentiguide-Puro as described by the authors to generate the pLentiguide-Puro-TMEM70_sgRNA-1 to 4. plentiCas9-Blast and lentiGuide-Puro were gifts from Feng Zhang (Addgene # 52962 and # 52963 plasmids). The sequences of all primers used for these constructs can be found in Table 1.

**Table 1.**
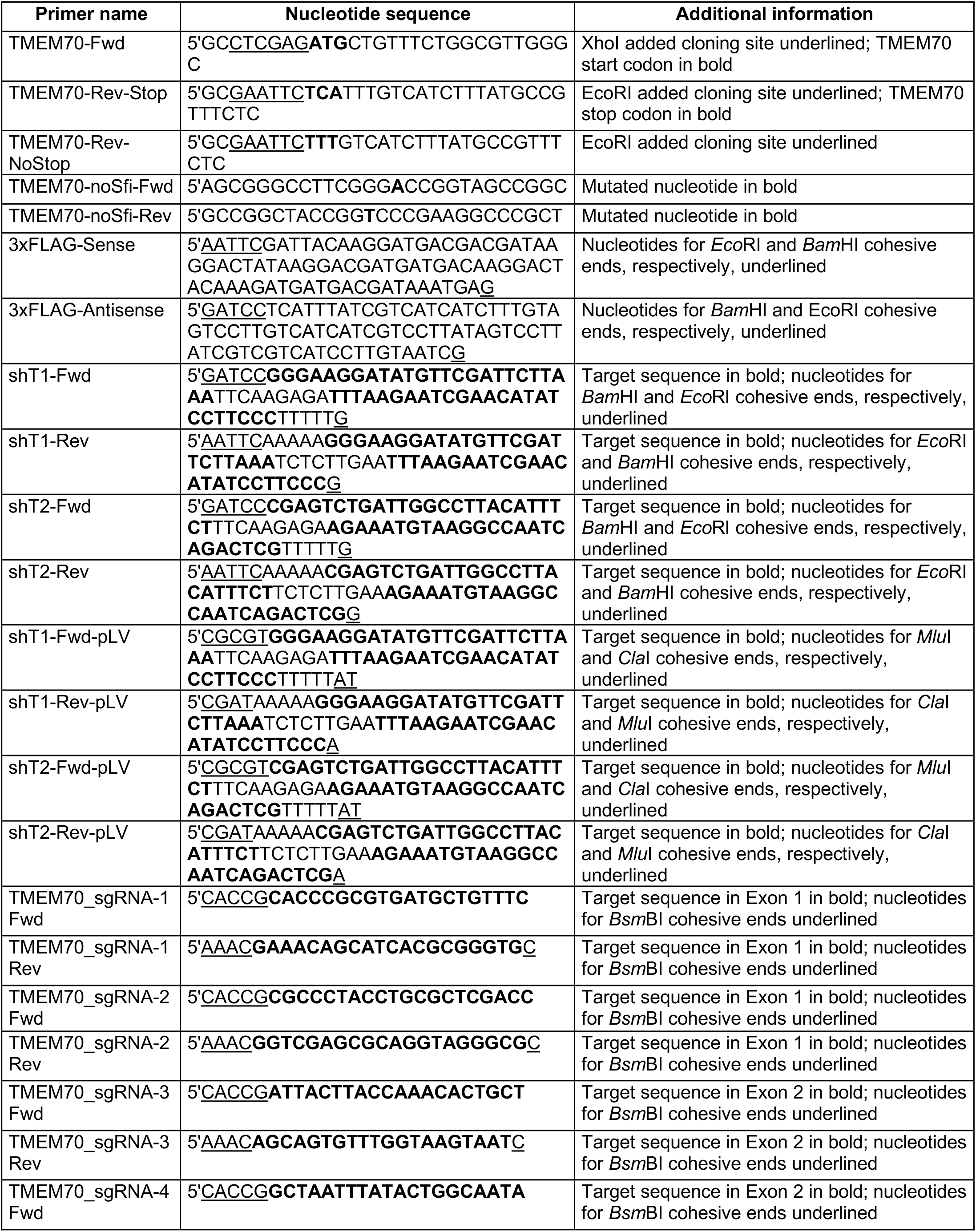
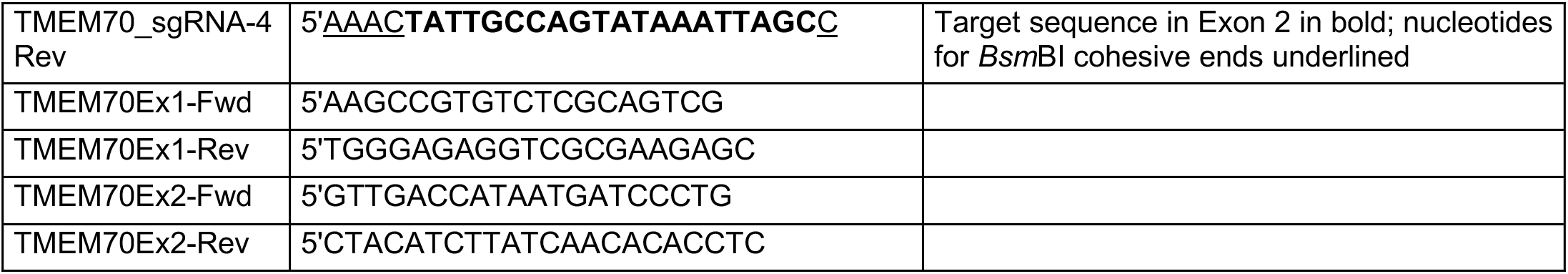
Oligonucleotides used in this study.

### Generation of TMEM70 KO cell lines by CRISPR-Cas9

HEK293T cells were co-transfected with pLentiCas9-Blast and either of the sgRNA constructs targeting the TMEM70 gene. 48 hours after transfection, the cells were passaged and subjected to antibiotic selection (Blasticidin 20 μg/ml and Puromycin 2 μg/ml) for three days. Cells were then allowed to recover and amplify without antibiotics for 5 days before samples were taken for analysis and frozen stocks of viable cells were prepared. sgRNA-2 and sgRNA-3 series were the most effective constructs, *i.e.* with minimal signal of TMEM70 protein by western blot. Corresponding cells were cultured again and cloned in 96 well-plates by critical dilution. A first screening by western blot allowed to select clones showing no expression of TMEM70 of which three from each series were selected (T2AC2/T2AC11/T2BB9 for sgRNA-2, and T3AA2/T3AG7/T3BH6 for sgRNA-3). For these clones, the exon (n°1 or 2) of TMEM70 targeted by the corresponding sgRNA was amplified by PCR using the primer pairs TMEM70Ex1-Fwd/TMEM70Ex1-Rev or TMEM70Ex2-Fwd/TMEM70Ex2-Rev, respectively, and sequenced in both directions using the same primers. The chromatograms were analyzed using the ICE algorithm (https://ice.synthego.com/#/) that revealed most CRISPR edits. When required, the PCR products were cloned into the pGEM-T vector (Promega). Sequencing of the inserts from a dozen of subclones in each case allowed us to ascertain the edits in TMEM70 exons. All edits were short deletions/insertions close to the expected cut site leading to frame-shifts and early stop codons.

### Generation of stable cell lines

Lentiviral particles were generated in HEK293T cells according to (69). Cell lines were transduced with lentiviruses at a multiplicity of infection (MOI) of 1 unless otherwise specified and incubated for two to three days. Transduced cells were further amplified and either selected using antibiotic resistance (Su.Beta knockdown) or subjected to fluorescence assisted cell sorting (FACS) according to the fluorescent proteins co-expressed by the plasmids. For RNA interference studies the same rationale was used with additional peculiarities. In order to achieve doxycycline inducible TMEM70 knock-down, HEK293T cells were first transduced with lentiviruses bearing the pLV-tTR-KRAB-red construct at MOI 2 and were subjected to a second transduction with both the pLVTHM-T1 and pLVTHM-T2 constructs at MOI 5 each. shRNAs induction was triggered by addition of doxycycline 250 ng/ml. Cells were cultured in the presence of doxycycline for various times with medium replacement every two to three days. To perform the Su.Beta knockdown, HEK293T were transduced with TRC2-pLKO-puro-shBeta3 and TRC2-pLKO-puro-shBeta4 lentiviral particles at MOI 3 each. For TMEM70 and Su.Beta double knock-down, the same cells were further transduced with pSIH-T1 and pSIH-T2 at MOI 3 each.

### Primary Antibodies

Anti-TMEM70 antibodies were rabbit polyclonal sera raised by Eurogentec SA (Belgium) against a mixture of two peptides: an internal peptide located within the N-terminal part of the mature protein TPSDKSEDGRLIYTG and the C-terminal peptide YMEETSEEKRHKDDK, respectively P1 and P2 in Fig. 1A. TMEM70 anti-sera were used at a 1:1000 dilution in western blots. Reactivity of sera against each peptide was validated by ELISA (See Fig. S1A).

The following commercial primary antibodies were used throughout this study: Mouse monoclonal antibodies against Actin (clone C4 MAB1501R, Millipore), Fibrillarin (ab18380, Abcam), Cytochrome c (556433, BD Biosciences), VDAC1 (ab14734, Abcam), anti-FLAG M2 (F1804), Core2/UQCRC2 (ab14745, Abcam), NDUFB6 (ab110244, Abcam), SDHA (ab14715, Abcam), Su.Alpha/ATP5A (ab14748, Abcam), Su.Beta/ATPB (ab14730, Abcam), Su.d/ATP5H (ab110275, Abcam), Su.b/ATP5F1 (ab117991, Abcam), Su.OSCP/ATP5O (ab110276, Abcam); Rabbit Polyclonal antibodies against ERGIC p58 (E1031, Sigma), Mitofilin/Mic60 (10179-1-AP, Proteintech), TOM20 (SC-11415, Santa-Cruz); and Rabbit monoclonal antibodies against Su.c/ATP5G1/G2/G3 (ab180149, Abcam).

### Subcellular fractionation

Cell fractionation protocol was adapted from previously established protocols (70). Cells were harvested, washed in PBS, resuspended in a HEPES 10 mM buffer at pH7.4 containing Mannitol 210 mM, Sucrose 70 mM, EDTA 1 mM, cOmplete^TM^ protease inhibitor cocktail (MB buffer) and homogenized on ice by several passages through a 25G needle fitted to a 2-5 ml syringe. All further steps were done at 4°C or on ice. The cell extract was first centrifuged at 700g for 10 min in order to pellet nuclei, cell debris, and unbroken cells. These steps were repeated until almost no unbroken cells were remaining in the pellet. Supernatants were further centrifuged at 1.000 g and all pellets were pooled in the nuclei enriched fraction. Supernatants from the last step were pooled and further centrifuged at 10.000 g for 10 min to pellet a fraction enriched in mitochondria (corresponding to the isolated mitochondria used throughout this study). Last supernatant was then subjected to ultracentrifugation at 130.000 g for 30 min in order to pellet a microsomal fraction and isolate a cytosolic fraction in the supernatant. Proteins from all factions were precipitated with trichloroacetic acid, centrifuged at 25.000 g for 10 min, washed with cold acetone, allowed to dry and solubilized in loading buffer.

### Submitochondrial fractionation

Salt and carbonate extraction were adapted from previously established protocols (70). Briefly, isolated mitochondria were resuspended in HEPES-KOH 20 mM buffer supplemented with either KCl 500 mM at pH7.4 and sonicated 10 times for 10 seconds with 10 seconds breaks, or Na_2_CO_3_ 0.1 M at pH11 and sonicated once 10 seconds and incubated for 30 min on ice. All samples were centrifuged for 10 min at 16.000 g. All supernatants and pellets were treated as described for cell fractions. For protease protection assay, isolated mitochondria were either resuspended in MB buffer or swollen by resuspension in HEPES-KOH 20 mM pH7.4. For each buffer condition, two tubes were prepared among which one was supplemented with 200 μg/ml Proteinase K. After incubating for 30 min on ice, 2 mM PMSF were added to samples to inhibit the proteinase K and further incubated for 5 min on ice. All samples were centrifuged for 10 min at 16.000 g. Pellets were treated as described for cell fractions.

### Total protein extracts

Cells were harvested, washed with PBS, resuspended in 10 volumes of RIPA buffer (Tris-HCl 50 mM pH8.0, 150 mM NaCl, 1% IGEPAL®CA-630, 0.5% sodium deoxycholate, 0.1% sodium dodecyl sulfate) supplemented with cOmplete™ protease inhibitor cocktail (Roche) and incubated for 20 minutes on ice. The suspensions were then centrifuged at 4°C for 20 minutes at 12000g and supernatants were transferred to fresh tubes. Protein concentrations were determined using the BCA assay kit (Pierce).

### SDS-PAGE and western blot

Protein samples were lysed in loading buffer (Tris-HCl pH6.8 62.5 mM, Glycerol 10%, SDS 2%, bromophenol blue 0.002%, dithiothreitol 100 mM) and separated by SDS-PAGE using NuPAGE Bis-tris or Bolt Bis-Tris (12% or 4-12%) precast gels (Invitrogen) and MOPS or MES running buffers, respectively. Proteins were then transferred to nitrocellulose membrane in Tris 25 mM, Glycine 192 mM, 20% ethanol using a Mini Trans-Blot® cell (Bio-Rad), probed with desired primary antibodies in TBS-T buffer (Tris 20 mM pH8.0, NaCl 137 mM, 0.05% (w/v) Tween-20), supplemented with 5% (w/v) non-fat dry milk, revealed by chemiluminescence using peroxidase-conjugated Goat anti-Rabbit or Goat anti-Mouse IgG (H+L) secondary antibodies (Jackson ImmunoResearch) and Clarity™ western blot ECL substrate (BioRad). Signals were acquired using a G:BOX (Syngene) and quantified using the ImageJ software.

### BN-PAGE of isolated mitochondria

BN-PAGE experiments were performed as described previously (71, 72) with some modifications. Mitochondria (300 μg mitochondrial proteins) were resuspended in 30 μl of 30 mM HEPES pH7.4, 150 mM potassium acetate, 12% (w/v) glycerol, 2 mM 6-aminocaproic acid, 1 mM EDTA, cOmplete^TM^ protease inhibitors (Roche), with digitonin (digitonin to protein ratio of 2 g/g), and incubated for 30 min at 4°c in a thermomixer (1400 rpm). Insoluble material was eliminated (pellet) by centrifugation at 4°C for 20 min at 25.000 g. 1μl of loading buffer (750 mM 6-aminocaproic acid 5% (w/v) Coomassie G-250) was added to the protein extract (supernatant) before loading on the gel. BN-PAGE was performed using NativePAGE 3-12% or 4-16% Bis-Tris gels and associated running buffer (Invitrogen). The cathode buffer was supplemented with 0.002% (w/v) Coomassie G-250. After electrophoresis, gels were either incubated in a solution of Lead acetate 0.5 g/L, ATP 4 mM, Glycine 270 mM, MgSO4 14 mM, Triton X-100 0.1%, Tris 35 mM pH8.4 to detect ATPase activity or subjected to two-dimensional SDS-PAGE.

### Two-dimensional electrophoresis BN-PAGE/SDS-PAGE

To perform two-dimensional electrophoresis, we used 2D NuPAGE gels and adapted protocols from the manufacturer’s instructions and previous reports (73, 74). Individual lanes from the BN-PAGE were excised and first equilibrated for 5 min in Tris 50 mM pH8.5, urea 6M, glycerol 30%, SDS 2% at room temperature under gentle agitation, then for 20 min in reducing conditions in the same buffer supplemented with DTT 100 mM, followed by a 20 min alkylation step in equilibration buffer containing iodoacetamide 135 mM, and a final 5 min washing step in 2D equilibration buffer. The gel lane was then transferred to a 2D well of a NuPAGE Bis-Tris 4-12% gradient gel. After the migration, the proteins were transferred to nitrocellulose as described for SDS-PAGE and analyzed by western blot.

### Immunoprecipitation

Mitochondrial extracts were prepared exactly as described for BN-PAGE, then 10-fold diluted in TBS buffer (50mM Tris-HCl pH 7.5; 150mM NaCl) supplemented with cOmplete^TM^ protease inhibitors (Roche). ANTI-FLAG M2 Magnetic Beads (SIGMA) were prepared as described by the manufacturer, resuspended in the end in TBS, mixed with approximately equal volume of mitochondrial lysate, and incubated for 2 hours at room temperature in an end over end shaker. The supernatant (flow-through) was aspirated and the beads were then washed three times with TBS buffer supplemented with 0,1% digitonin and eluted by competition with 150μg/ml 3xFLAG peptide (Sigma) in TBS + 0,1% digitonin for 30 minutes at 4°C in an end over end shaker.

### Immunofluorescence

Human skin fibroblasts or 143B cells expressing TMEM70-FLAG were seeded at a density of ~100,000 cells per well in a six-well plate containing 12-mm coverslips and incubated overnight. Cells were fixed with 4% paraformaldehyde (PFA) in PBS for 20 min. After fixation, cells were quickly washed 3 times in PBS and post-extracted for 10 min in PBS with 0.1% Triton X-100. PFA free radicals in excess were reduced with 10 mM NH_4_Cl in PBS for 10 min, then cells were incubated in the blocking buffer (BF; PBS with 1% BSA) for 15 min. Next, coverslips were incubated with primary antibody diluted in the BF for 1h at 37°C. Coverslips were then washed in BF three times for 5 min and subsequently incubated for 45 min at 37°C with secondary antibody diluted in BF in dark chamber. Finally, coverslips were washed in PBS three times for 5 min, and mounted on a glass slide with ProlonGold® mounting medium. For immunolabeling, primary antibodies were rabbit polyclonal anti-TMEM70 (1:100), rabbit polyclonal anti-MIC60 (1:200), mouse monoclonal anti-FLAG (1:500); and secondary antibodies were goat anti-mouse Alexa 647 IgG (H+L), goat anti-rabbit Alexa 568 IgG (H+L) (1:100; Invitrogen), goat anti-mouse Atto 647N (1:100; Sigma-Aldrich), or anti-rabbit STAR-580 and anti-mouse STAR-RED (1:200; Abberior) for STED acquisition. For Expansion microscopy, after the last PBS wash of standard immunofluorescence, cells were proceeded exactly as described by (46).

### Optical Microscopy

Expanded cells were imaged with a UPlanSApo 100×/1.4 oil immersion objective in an Olympus IX81 microscope (Olympus, Tokyo, Japan). Z-stacks were deconvolved using the DeconvolutionLab plugin. Stimulated Emission Depletion (STED) imaging was performed using a STEDYCON (Abberior Instruments GmbH, Göttingen) with excitation lasers at 488, 561, and 640 nm, and a STED laser at 775 nm (maximum intensity 1.25 W; all lasers are pulsed with 40 MHz repetition rate). The STEDYCON was mounted on the camera port of an AxioImager Z2 upright microscope (Zeiss, Jena, Germany), equipped with a 100x objective (alpha Plan-Apochromat, Oil, DIC, Vis, NA 1.46; Zeiss). The pinhole was set to 1.1 Airy units for 650 nm emission. Fluorescence was detected on avalanche photo diodes, with emission bands between 650–700 nm, 578–627 nm, and 505–545 nm, respectively. Data was stored in .obf format and exported as tif files for further analysis.

### Electron microscopy

Cells were fixed for 45 min at 4 °C in 4% PFA in PBS, washed and fixed again for 1 h at room temperature in 2% osmium tetroxide in PBS containing 15 mg/ml of K_4_Fe(CN)_6_. Dehydration was performed with ethanol series (50%, 70%, 95% and absolute ethanol). Thereafter, the samples were embedded progressively in Epon resin. Ultrathin sections were contrasted with 1% lead citrate in water for 1 min. Sample imaging was performed using a Hitachi H7650 transmission microscope operated at 80 KV with a Gatan—11 MPx camera at electron microscopy unit (Bordeaux imaging center).

### Oxygen consumption measurements

Cells from around 60% confluent Petri dishes were detached using trypsin, harvested by centrifugation at 1.000g for 8 min at 4°C and resuspended in 100 μl PBS. Oxygen consumption rates were measured with 1.10^6^ cells/ml by high resolution respirometry in an Oxygraph-2k (Oroboros Instruments) in respiration buffer (sucrose 120 mM, KCl 50 mM, KH_2_PO_4_ 4 mM, MgCl_2_ 2 mM, EGTA 1 mM, Tris-HCl 20 mM pH7.2) supplemented subsequently with succinate 5 mM, ADP 2.5 mM, oligomycin 2 μg/ml, and CCCP 0.25 μM.

## Data Availability

All the data are contained within the manuscript and supplemental information.

## Acknowledgements

We are grateful to the Genetic unit of Hôpital Necker-Enfants malades, Paris, France and Agnès Roetig (Institut Imagine, UMR1163, Paris) for providing primary skin fibroblasts from a TMEM70 patient and a control individual. We thank Anne Chomyn, Feng Zhang, and Didier Trono for providing tools and reagents; Michael Zick for the cloning of pKSPS-FLAG; Claire Dubos and Maurean Guerreiro for technical assistance. We thank the members of the Devin lab and Marie-France Giraud, Alain Dautant, and François Godard for fruitful discussions during the preparation of this manuscript. STED microscopy was performed using a STEDYCON kindly lent by Abberior Instruments GmbH, Göttingen. Electron microscopy imaging was performed at the Bordeaux Imaging Center. We thank Vincent Pitard for technical assistance at the Flow cytometry facility (CNRS UMS 3427, INSERM US 005, Univ. Bordeaux), and Véronique Guyonnet-Dupérat (Vect’UB vectorology facility) for helpful discussions and technical advice in the knockdown experiments. This work was supported by the CNRS (Centre National de la Recherche Scientifique). HB was funded by the Tunisian ministry for higher education and scientific research.

## Author contributions

SDC, AD, and JPdR, and ET conceived the study. HB, JB, and SDC designed and conducted the majority of experiments and analyzed the data. JD and MR designed and performed immunofluorescence microscopy experiments and data analysis. BS and CB performed electron microscopy experiments and data analysis. MRose designed, performed and analyzed part of knockdown and 2D experiments. SC provided technical support in CRISPR/Cas9 experiments. SDC and ABAE supervised the project. SDC wrote the original draft and reviewed and edited the manuscript together with AD, JPdR, MR, and ET. All authors approved the final version of the manuscript.

