## Supplementary Figures for "TMEM70 forms oligomeric scaffolds within mitochondrial cristae promoting *in situ* assembly of mammalian ATP synthase proton channel"

- Supplementary Figures:
  - Figure S1
  - Figure S2
  - Figure S3
  - Figure S4
  - Figure S5
  - Figure S6

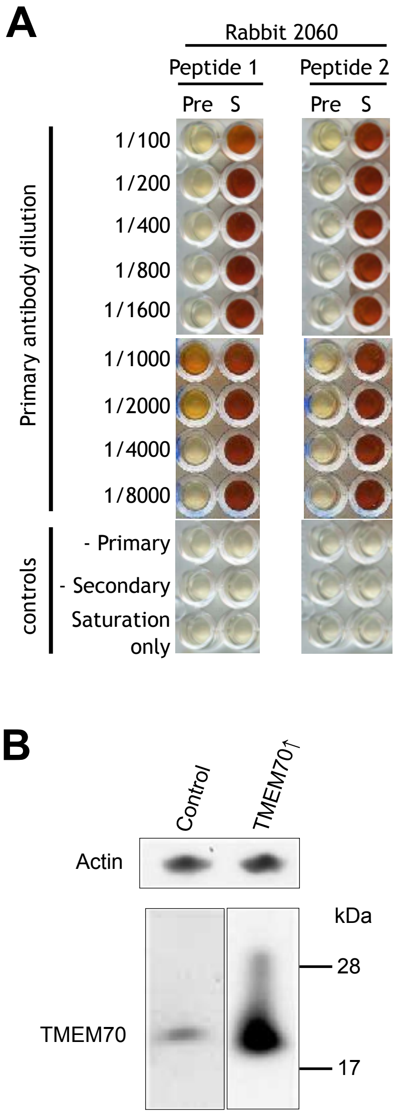


**Figure S1. TMEM70 antibodies validation.** *A*, ELISA test of anti-TMEM70 serum. The serum was tested separately against the two immunogenic peptides. Each peptide was bound to the surface of the wells. The preimmune (Pre) or immune (S) serum from rabbit 2060 was applied to the wells after dilution 1:100 to 1:8000 fold as indicated on the figure. After incubation with peroxidase-conjugated anti-rabbit antibodies, the relative amount of primary antibodies bound to the peptides was assessed by color intensity using OPD substrate. Additional controls were performed omitting primary sera (secondary), secondary antibodies (primary), or both (saturation). *B*, total protein extracts from 143B-rho$^{+}$ cell line overexpressing (TMEM70↑) or not (Control) were analyzed by SDS-PAGE and western-blot using antibodies against indicated proteins.

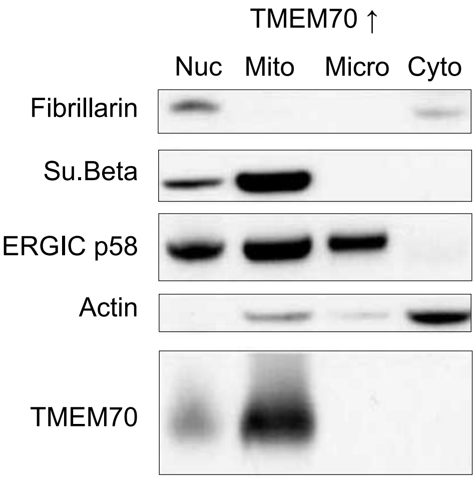


**Figure S2. Subcellular localization of exogenous TMEM70.** 143B-rho$^+$ cells overexpressing TMEM70 (TMEM70↑) were fractionated by differential centrifugation as detailed in the materials and methods section. Fractions enriched in nuclei (Nuc), mitochondria (Mito), microsomes (Micro), and cytosol were analyzed by SDS-PAGE and western blot using antibodies against indicated proteins.

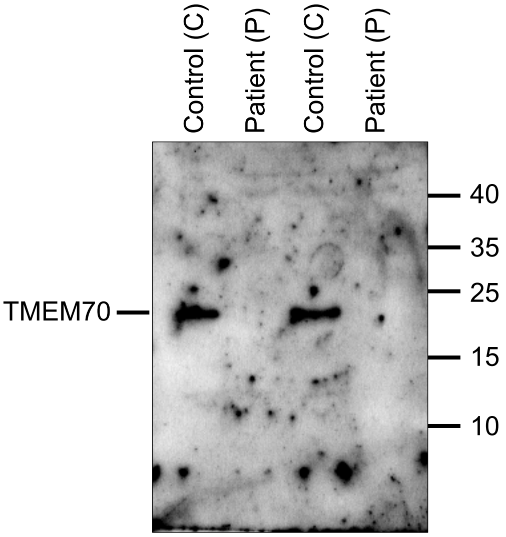


**Figure S3. Characterization of TMEM70 in patient skin fibroblasts bearing the c. 497_498del mutation in TMEM70.** Isolated mitochondria from cultured fibroblasts of the patient and a control individual were lysed and proteins were separated by SDS-PAGE and analyzed by western blot using anti-TMEM70 serum. The figure shows an extended view of the western blot result presented Fig. 4B together with the positions and sizes of the molecular markers in KDa.

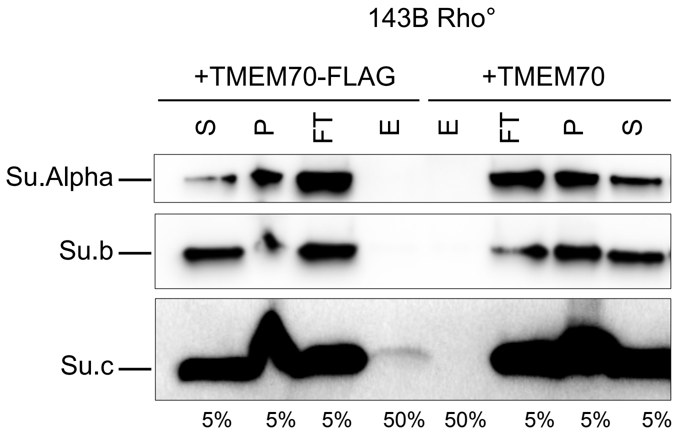


**Figure S4. Physical interaction between TMEM70 and the subunit c of the ATP synthase in Rho° cells.** Immunoprecipitation of TMEM70-FLAG. Mitochondria isolated from Rho° cells overexpressing TMEM70-FLAG, or TMEM70 as a control, were used in immunoprecipitation assays using anti-FLAG M2 magnetic beads. Digitonin extractions were performed in order to preserve protein-protein interactions. Extraction supernatants (S) represent the input material incubated with the beads while insoluble pellet (P) were kept apart for analysis. The unbound material come with the flow-through (FT) while bound proteins are retained on the beads and further eluted (E) natively by competition with free 3xFLAG peptide. Portions (indicated at the bottom of the panel) of each fraction were analyzed by SDS-PAGE and western blot.

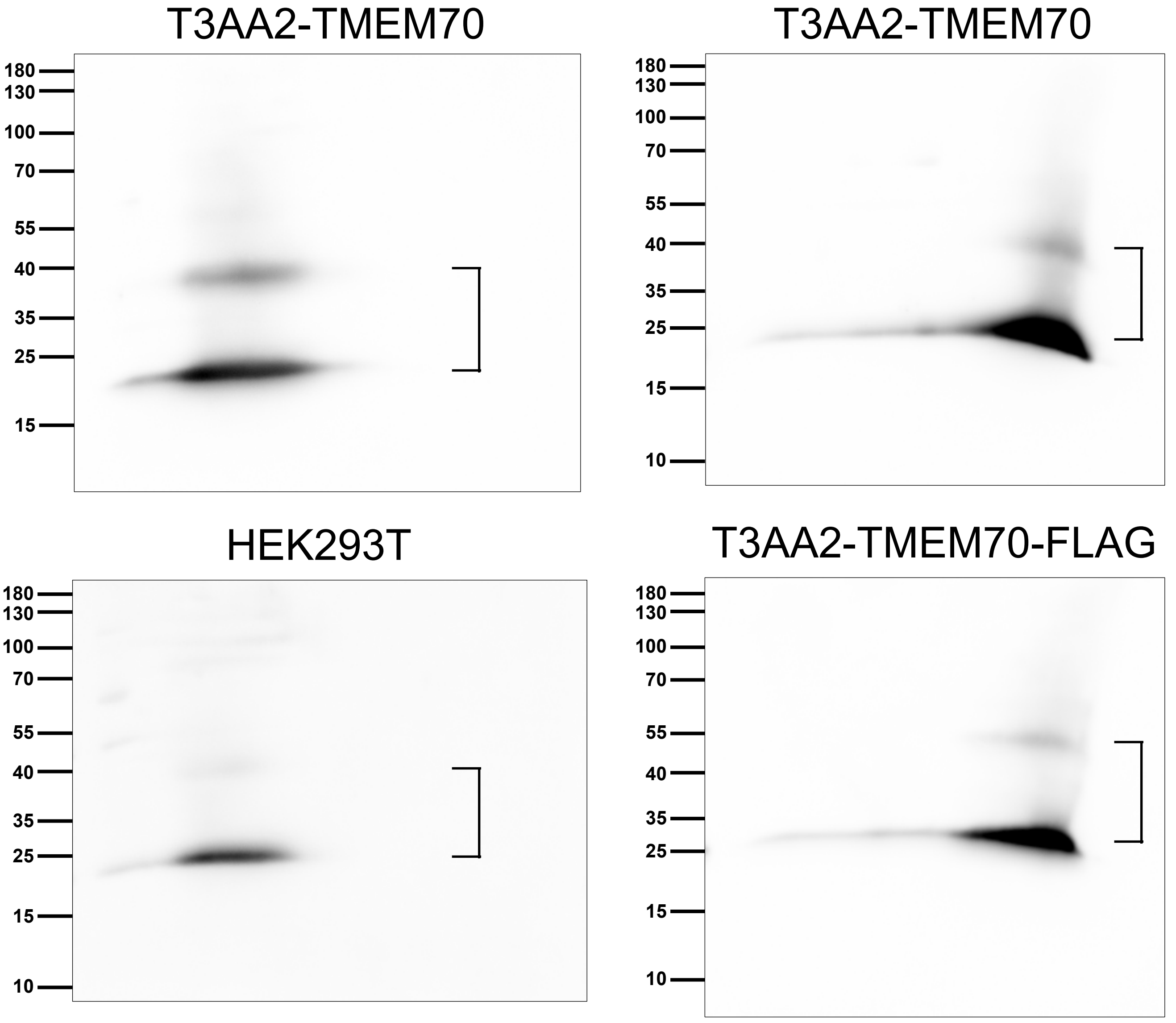


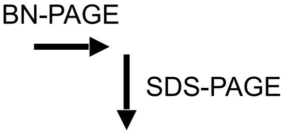


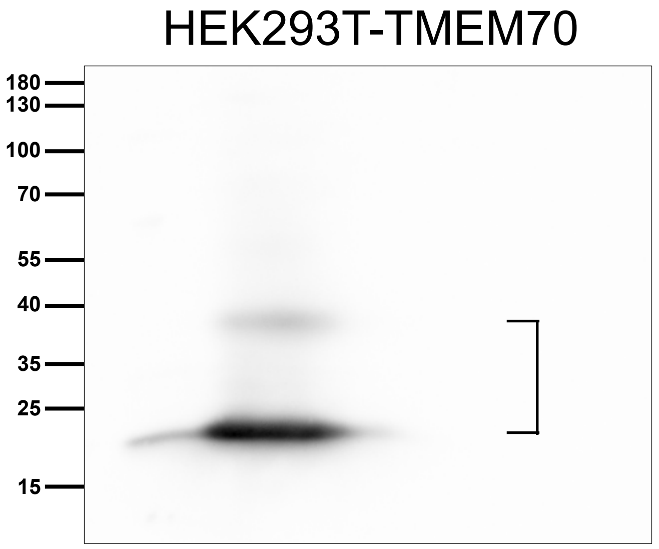


**Figure S5. Visualization of TMEM70 dimers in SDS-PAGE second dimensions after a first Blue Native PAGE.** Mitochondria were isolated from indicated cell lines and subjected to digitonin extraction. The extracts were analyzed by BN-PAGE in a first dimension. Lanes of the BN-PAGE were cut and analyzed by SDS-PAGE in a second dimension. Proteins were transferred to nitrocellulose and analyzed by western blot. Arrows indicate the migration direction for the two dimensions. The figure shows an extended view of the western blot result presented Fig. 6 with antibodies directed against FLAG or TMEM70, together with the positions and sizes of the molecular markers in KDa. The position of the signals for TMEM70 monomers and dimers are indicated.

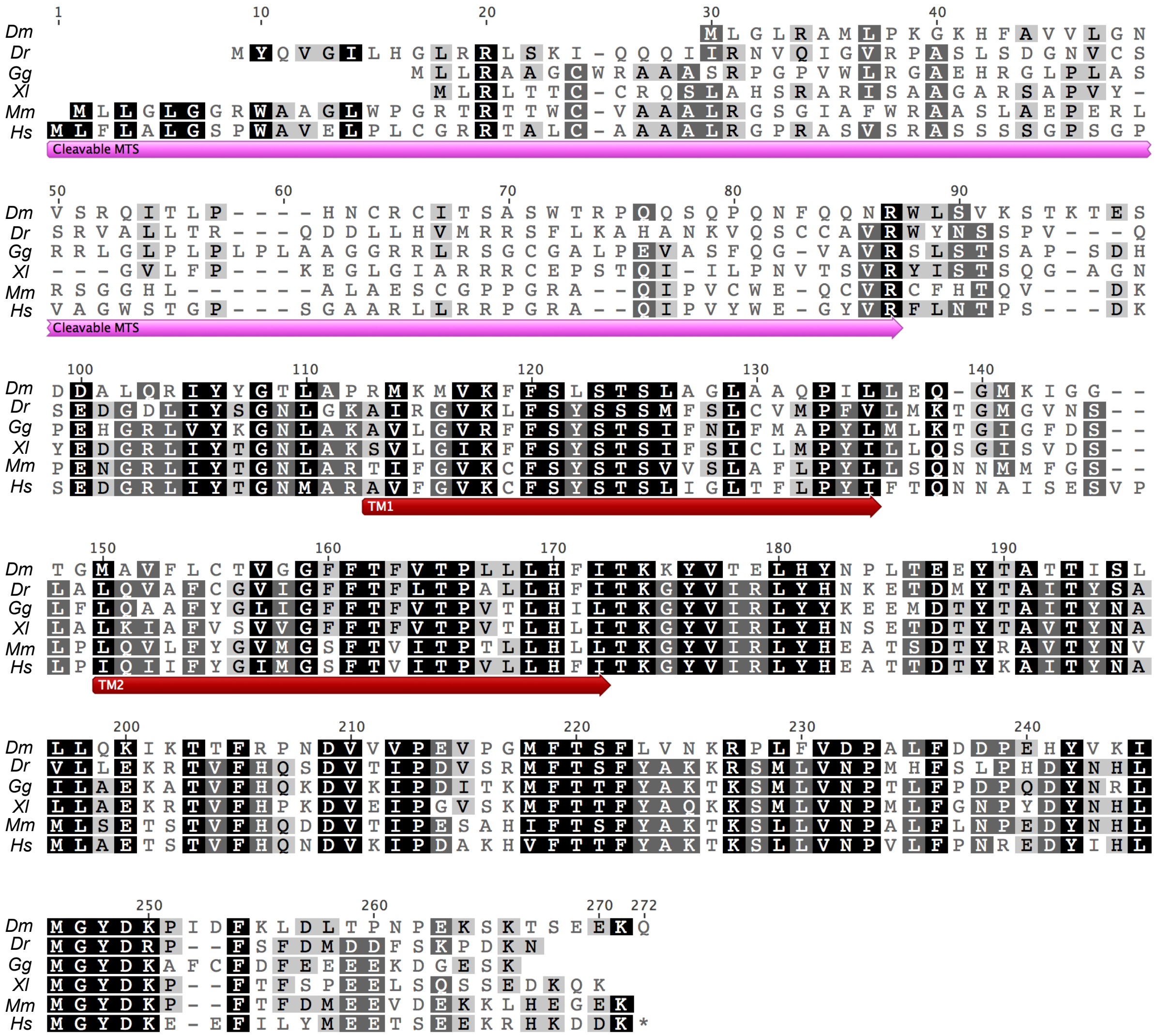


**Figure S6. Alignment of TMEM70 primary sequences from different species.** Primary sequences of TMEM70 precursors from indicated species were aligned using the ClustalW2 algorithm. Conserved residues are highlighted using a grey range from identical or very similar residues in all species in black to non-conserved residues in white. Predicted mitochondrial targeting sequence (cleavable MTS) and positions of the predicted transmembrane segments (TM1, TM2) of the human protein are indicated in pink and red, respectively. Drosophila melanogaster Q95SS8 (Dm); Danio rerio F1Q9S8 (Dr); Gallus gallus Q5ZLJ4 (Gg); Xenopus laevis A0A1L8FT80 (Xl); Mus musculus Q921N7 (Mm); Homo sapiens Q9BUB7 (Hs).
